# Deep and superficial layers of the primary somatosensory cortex are critical for whisker-based texture discrimination in mice

**DOI:** 10.1101/2020.08.12.245381

**Authors:** Jung M Park, Y Kate Hong, Chris C Rodgers, Jacob B Dahan, Nina Harano, Ewoud RE Schmidt, Randy M Bruno

## Abstract

The neocortex, comprised of multiple distinct layers, processes sensory input from the periphery, makes decisions, and executes actions. Despite extensive investigation of cortical anatomy and physiology, the contributions of different cortical layers to sensory guided behaviors remain unknown. Here, we developed a two-alternative forced choice (2AFC) paradigm in which head-fixed mice use a single whisker to either discriminate textures of parametrically varied roughness or detect the same textured surfaces. Lesioning the barrel cortex revealed that 2AFC texture discrimination, but not detection, was cortex-dependent. Paralyzing the whisker pad had little effect on performance, demonstrating that passive can rival active perception and cortical dependence is not movement-related. Transgenic Cre lines were used to target inhibitory opsins to excitatory cortical neurons of specific layers for selective perturbations. Both deep and superficial layers were critical for texture discrimination. We conclude that even basic cortical computations require coordinated transformation of sensory information across layers.

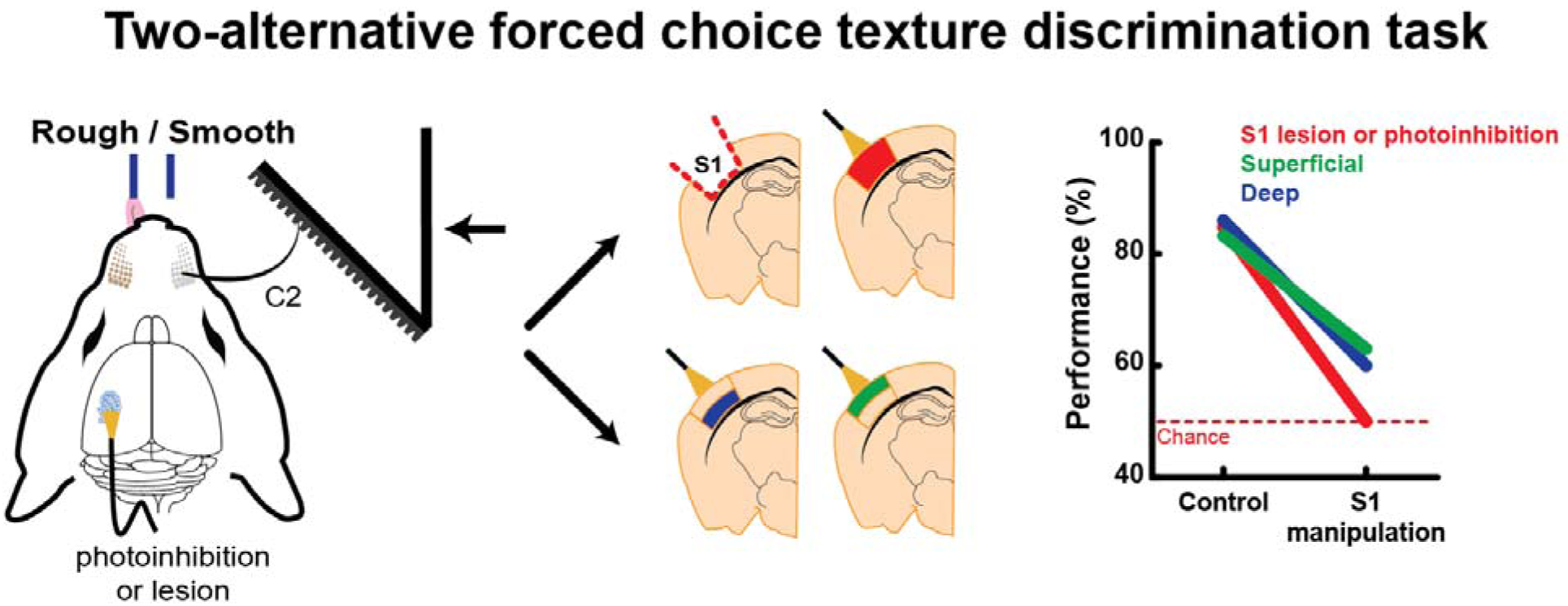

## Introduction

As animals navigate through their environment, they must acquire sensory information and extract behaviorally relevant features to make effective decisions. While the cortex plays an important role in sensory perception, our understanding of how cortical microcircuits functionally contribute to sensory guided behaviors is limited (Adesnik & Naka 2018, Douglas & Martin 2004). One of the most prominent features of cortical microcircuitry is its highly conserved laminar organization. The differing cell types and response properties of each layer suggest distinct functional roles (Petersen 2019). Sensory information is known to arrive from thalamus in cortical layer 4 (L4), but sparse thalamocortical branches in L5/L6 can account fully for excitation in the deep layers of somatosensory cortex (Constantinople & Bruno 2013). Similar results have been demonstrated in visual cortex (Pluta et al 2015). This duplication of ascending sensory information suggests that the cortex contains two separate processing systems composed of deep and superficial layers that may function independent of one another.

Investigating the relationship of these cortical microcircuitry features to behavior requires a cortex-dependent task. Whisker-mediated tactile behaviors are ideal for this purpose because (1) the one-to-one correspondence of whiskers and columns in the rodent barrel cortex, a subregion of primary somatosensory cortex (S1), allows precise targeting of relevant microcircuits and (2) genetic tools for manipulating neuronal activity are readily available in the mouse (O’Connor et al 2009). Research into many sensory modalities, especially somatosensation, has probed cortical circuitry and function using detection paradigms, in which animals must report the presence or absence of a stimulus (Brecht et al 1997, Hutson & Masterton 1986). Neurons in S1 have activity that correlates with touch, and manipulating their activity can alter detection behavior (Guo et al 2014b, Miyashita & Feldman 2013, O’Connor et al 2010, Sachidhanandam et al 2013, Yu et al 2016). Yet we recently showed that barrel cortex is, surprisingly, dispensable for both performing and learning a whisker-based go/no-go object detection task (Hong et al 2018). A prominent role of cortex may materialize during more complex tasks.

Sensory discrimination is more computationally demanding than simple object detection. For example, texture discrimination requires the integration of tactile information over space and time (Lederman 1983, Weber et al 2013). Human stroke studies and classical lesion studies in nonhuman primates suggest that somatosensory cortex is essential to texture behaviors (Corkin et al 1970, Randolph & Semmes 1974). Rodents are highly adept at differentiating textures (Carvell & Simons 1990, Carvell & Simons 1995, Chen et al 2013, Diamond 2010). Only one study of the rodent whisker system has causally examined the role of cortex in texture discrimination, finding the cortex to be indispensable (Guic-Robles et al 1992). Those rats were given extensive post-lesion recovery periods prior to retesting. However, early retesting is critical to assess an animal’s ability to recover from lesion of either the somatosensory or motor systems, whether the task is cortex independent or dependent (Hong et al 2018, Ng et al 2015, Zeiler & Krakauer 2013).

Building on these studies, we set out to develop a more sophisticated texture discrimination paradigm that facilitates the functional study of cortical layers. A design with two overt choices would avoid confounds of lapses in animal engagement found in common go/no-go designs (Shenoy & Angela 2012). Ideally, textures could be parametrically varied to ensure complete control of spatiotemporal surface features that determine how rough surfaces *feel* to the animals (Cascio & Sathian 2001, Lederman 1983) as opposed to frequently used and highly variable sandpaper discriminanda. Additionally, a head-fixed setup would permit precise presentation of surfaces and constrain mice to use their whiskers (Guo et al 2014a). Finally, an ideal behavior would require only a single whisker, minimizing the extent of cortex that needs to be manipulated.

To that end, we developed a novel texture discrimination task to interrogate the functional contribution of cortical layers. We trained head-fixed mice to discriminate parametrically controlled surfaces in a two-alternative forced choice (2AFC) paradigm. The whisker pad was paralyzed to investigate if active sensation enhanced behavior. We lesioned trained mice to test whether texture discrimination requires barrel cortex. To determine whether the 2AFC design itself required cortex, we compared lesioned mice trained on a 2AFC object detection task that used the same textured surfaces. Using targeted optogenetic inactivation of cortical layers, we investigated the functional role of both the deep and superficial layers in texture discrimination.

## Results

### Parametric texture discrimination tasks

We developed a novel 2AFC texture discrimination paradigm for head-fixed mice. At the onset of each trial, a linear actuator advanced a textured surface into the mouse’s right whisker field, 45° relative to the animal’s midline (Fig 1A, Video 1). In this version of the task, textures were passively presented (so close to the face that active whisking was not required to sample them). Even in the absence of whisking, most whiskers were in contact with the textures at the final stimulus position. Mice were trained to classify textures by licking one of two ports. A response window in which licks were counted as choices opened shortly after the surface reached its final position, but the mice often licked beforehand, cued to the trial initiation by the sound of the actuator.

**Figure 1.**
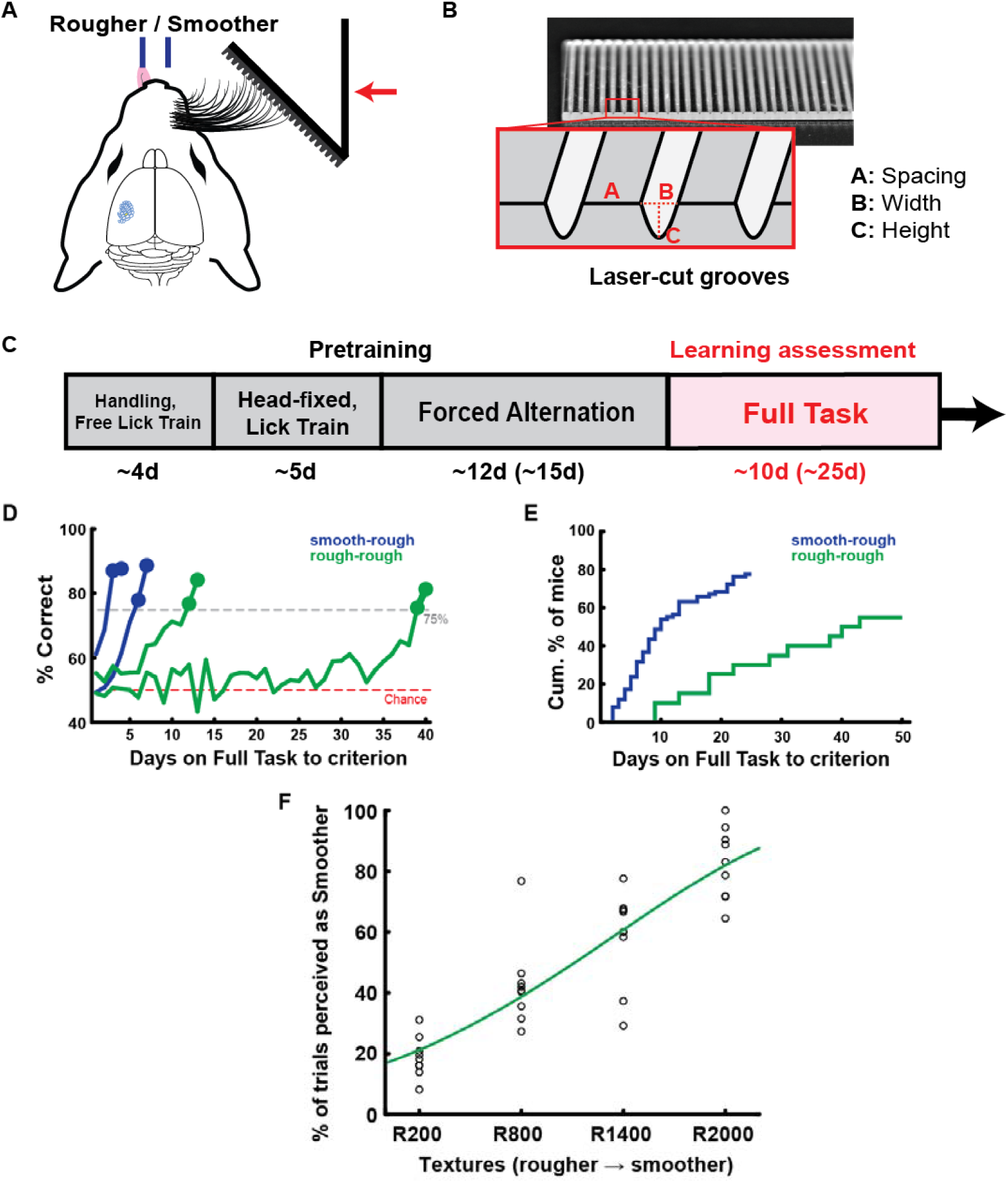
Mice readily discriminate textures in a two-alternative forced choice task. **(A)** Head-fixed two-alternative forced choice texture discrimination paradigm. Mice learned to lick left for rougher textures, and lick right for smoother textures. **(B)** Laser-cut surfaces allow for parametrically controlled textures. See Tables 1 and 2 for quantifications. **(C)** Training timeline for smooth-rough texture discrimination. Increased number of training days for rough-rough discrimination in parenthesis. **(D)** Example learning curves of mice reaching learning criterion (≥75% correct on two consecutive learning assessment sessions within 25 Full Task days for smooth-rough, 50 days for rough-rough). **(E)** Cumulative histogram of number of sessions to reach learning criterion (smooth-rough n = 59/76 mice, rough-rough n = 11/20). **(F)** Psychometric curve for first day on all four textures (n = 9 mice).

Our simplest version of the task involved a smooth-rough texture pair, for which the mice learned to lick right for a smooth surface and left for a grooved laser-cut surface (Fig 1B, Table 1) to obtain water rewards. Incorrect responses were punished with a 9-s timeout. After several pretraining phases to familiarize the mice with the 2AFC setup, mice began the Full Task phase in which trial types were randomized unless response biases were algorithmically detected (Fig 1C, see Methods). We quantified the number of days on the Full Task required to reach learning criterion (≥75% correct performance for 2 consecutive days). The mice quickly learned the task (9.5 ± 0.9 d, mean±SEM; range 2-24 d; Fig 1D). Trained mice generalized to two additional rough textures we tested (Table 1), performing equally well if their training rough texture was replaced with a different rough surface (Supplementary Fig 1).

**Table 1.** Textures used in the smooth-rough discrimination paradigm. Mice readily discriminated between the smooth and any of the three rough textures.

| Textures | [A] Spacing ($\mu\text{m}$) | [B] Width ($\mu\text{m}$) | [C] Height ($\mu\text{m}$) |
| --- | --- | --- | --- |
| Smooth | ∞ | 0 | 0 |
| Rough<sup>1</sup> | 0 | 350 | 500 |
| Rough<sup>2</sup> | 0 | 500 | 375 |
| Rough<sup>3</sup> | 375 | 850 | 460 |

**Table 2.** Textures used in the rough-rough discrimination paradigm. Mice discriminated between the R200 and R2000 textures. The intermediate R800 and R1400 textures were added to identify the psychometric curve for whisker-based texture discrimination.

| Textures | [A] Spacing ($\mu\text{m}$) | [B] Width ($\mu\text{m}$) | [C] Height ($\mu\text{m}$) |
| --- | --- | --- | --- |
| R200 | 200 | 350 | 500 |
| R800 | 800 | 350 | 500 |
| R1400 | 1400 | 350 | 500 |
| R2000 | 2000 | 350 | 500 |

In a variation of this paradigm, a cohort of mice was trained to discriminate between two grooved textures that are more similar to one another than smooth and rough: one with grooves spaced 200 um apart and another 2000 um apart (Supplementary Fig 2A; Table 2, R200 and R2000). Mice learned to discriminate the rough-rough texture pair more slowly (24.5 ± 3.7 d, mean±SEM) than the smooth-rough texture pair (*P*<0.001, two-sample *t-*test; Supplementary Fig 2B, Fig 1D), and fewer rough-rough mice (55%) reached learning criteria than smooth-rough mice did (78%; *P*=0.043, Chi-Square; Fig 1E). Once trained, rough-rough animals readily generalized to discriminate additional intermediate textures (R800 and R1400) in a more complex 4-surface version of the task (Supplementary Fig 2A,C, Table 2, see Methods). The psychometric curve on the first day of the 4-surface task strongly suggests that mice are truly sensing roughness as opposed to other sensory features associated with the different surfaces (Fig 1F). Interestingly, with additional training, animal performance on the intermediate textures shifted over time; performance shifts suggest changes in the mice’s strategy rather than in discriminatory thresholds (Supplementary Fig 2C). The remainder of our study focused on smooth-rough and rough-rough (R200-R2000) comparisons.

### Mice can discriminate textures using a single whisker

Ideally, to test the necessity of neural structures to behavior, we could limit trained mice to using few or one whisker, which would correspond to a small number of relevant cortical columns that would need to be manipulated. We therefore trimmed the right whiskers of smooth-rough trained mice to three and eventually a single C2 whisker (Fig 2A, Video 2). While trimming to three whiskers minimally impacted performance (*P*=0.19, paired *t*-test; Fig 2B, green), the drop was more precipitous when subsequently reduced to the single C2 whisker (*P*=0.014, paired *t*-test; purple), and more pronounced when transitioning from all whiskers to a single one (*P*<0.001, paired *t*-test; blue). Nevertheless, single-whisker performance in all cases eventually recovered to full-whisker levels on smooth-rough comparisons (see Fig 2D, pre-lesion days -2 and -1), typically within 7-10 days.

**Figure 2.**
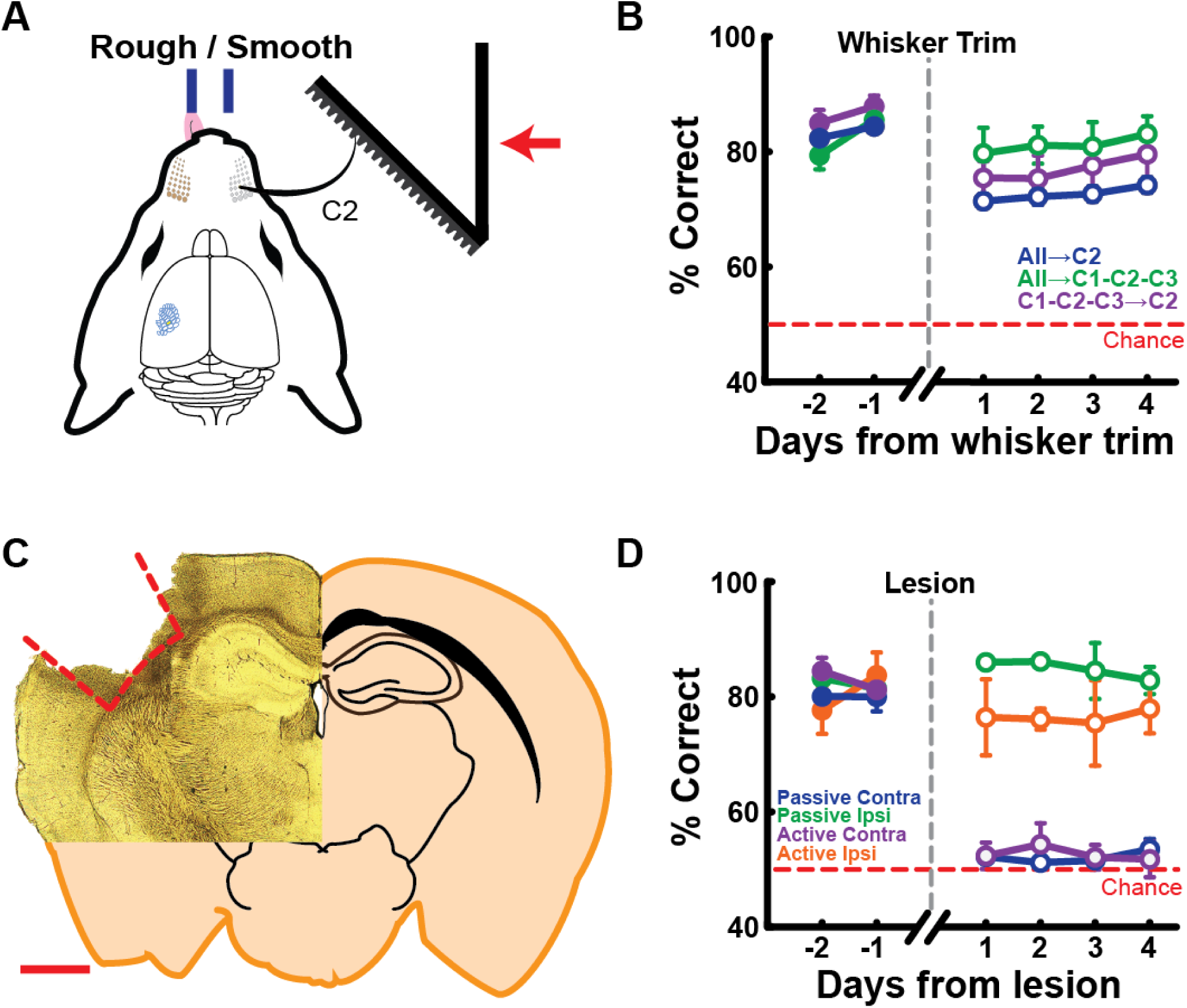
Texture discrimination can be performed with a single whisker and is cortex dependent. **(A)** Mice are trained to discriminate passively presented smooth-rough textures using their C2 whisker. **(B)** Mice can discriminate passively presented smooth-rough textures using a single whisker (all→C2 n = 39 mice, all→C1-C2-C3 n = 7 mice, C1-C2-C3→C2 n = 7 mice). Behavioral performance recovers to full-whisker levels (extended recovery beyond day 4 not shown). Mice can also learn to actively discriminate smooth-rough textures with a single whisker but require a period of additional training (learning curves not shown). **(C)** Coronal section of lesioned mouse brain. Red dashed lines, barrel cortex boundaries; scale bar, 1mm. **(D)** Behavioral performance in both passive and active versions of the task, before and after barrel cortex lesion (passive contralateral n = 10 mice, passive ipsilateral n = 3 mice, active contralateral n = 5 mice, active ipsilateral n = 3 mice). Data shown as mean±SEM.

Interestingly, mice were also able to discriminate between two grooved textures with three and even a single C2 whisker (Video 3). However, these rough-rough mice were strongly impaired by trimming, and their performance did not recover to full-whisker levels even after extended training (Supplementary Fig 2D), suggesting that finer discriminations require many whiskers. Thus, texture discrimination – whether smooth-rough or rough-rough – requires only a single whisker, but mice perform better by integrating information across whiskers when available. Because the single-whisker paradigm ensures better control for targeted manipulations, we trained animals to use a single whisker for manipulation experiments described below.

### Barrel cortex is indispensable for texture discrimination

Optogenetics and lesions can yield conflicting results (Hong et al 2018, Otchy et al 2015). Transient manipulations that result in behavioral effects may be due to diaschisis, in which other critical structures are inadvertently disrupted. Chronic manipulations can be more stringent tests of dispensability, assaying both short-term and long-term effects. Therefore, using intrinsic signal imaging, we targeted the cortical column corresponding to the remaining single whisker in smooth-rough mice and aspirated it and most of the surrounding columns (∼2 mm diameter, Fig 2C). Lesions that did not include layer 6 or that extended below the cortex into the striatum in post hoc histology were excluded from analysis. In mice passively presented with smooth-rough texture pairs, performance tested one day after lesioning the contralateral barrel cortex dropped to chance and showed no sign of recovery across successive days (*P*=0.002; paired *t*-test; Fig 2D, blue). By contrast, ipsilateral lesions, which controlled for non-specific effects of cortical damage and surgical procedure, had no effect on performance (*P*=0.45; paired *t*-test; green).

Active movements during sensation is thought to enhance discrimination behaviors (Gamzu & Ahissar 2001, Gibson 1962, Kleinfeld et al 2006). Active texture discrimination also differs from passive discrimination at a neural level by increasing activity in frontal and parietal cortex (Simões-Franklin et al 2011). We therefore trained an additional cohort to test whether barrel cortex is required for active texture discrimination. After initial training on the passive smooth-rough task, we gradually moved the textures further away until mice had to actively whisk forward to contact the textures with their remaining whisker (Video 4). Mice often required additional training (∼10-15 d) to master these new circumstances, but eventually could perform similarly to their purely passive smooth-rough counterparts (see pre-lesion Fig 2D). Similar to the passive conditions, the active smooth-rough cohort was severely impaired by barrel cortex lesion and showed no sign of recovery for 4 days (Fig 2D, purple), again in contrast to ipsilaterally lesioned mice, which were unaffected (orange). In summary, barrel cortex is indispensable for both passive and active 2AFC texture discrimination.

### Mice can discriminate textures without active movement

It is an open question whether active movement enhances texture discrimination capabilities in any species, including humans and non-human primates (Lederman 1981, Taylor & Lederman 1975). Contrary to the idea of movement enhancing discrimination, our active task animals, despite more extensive training, did not reach higher performance levels than passive task animals (see pre-lesion Fig 2D). However, high-speed videography showed passive task mice “pumping” their whisker against the passively presented textures in the final stimulus position (Video 2), perhaps acquiring information as freely moving rats exploring textures do when dragging their whiskers along surfaces (Carvell & Simons 1990, Carvell & Simons 1995, Jadhav et al 2009, Ritt et al 2008). It is therefore conceivable that mice convert the passive task into an active one. In the ideal comparison, the passive cohort should not be able to move its whiskers.

To address this, we paralyzed the muscles controlling the whisker pad by transecting the buccal and upper marginal motor nerves in single-whisker, passive smooth-rough trained animals (Fig 3A, Video 5). While there was a minor behavioral impairment the first day after the motor nerve cuts (*P*=0.014, paired *t*-test), the animals were able to discriminate smooth-rough textures without being able to move their whiskers (chance 50%; *P*<0.001, one-sample *t*-test; Fig 3B). By the second post-lesion session, the mice had recovered to full pre-lesion performance levels. This recovery did not involve mice switching from whisker-mediated touch to other sensory modalities, as whisker trimming reduced performance fully to chance (Fig 3C). Similarly, mice – albeit with all whiskers – could discriminate the more difficult rough-rough texture pair even after nerve cuts (Supplementary Fig 2E).

**Figure 3.**
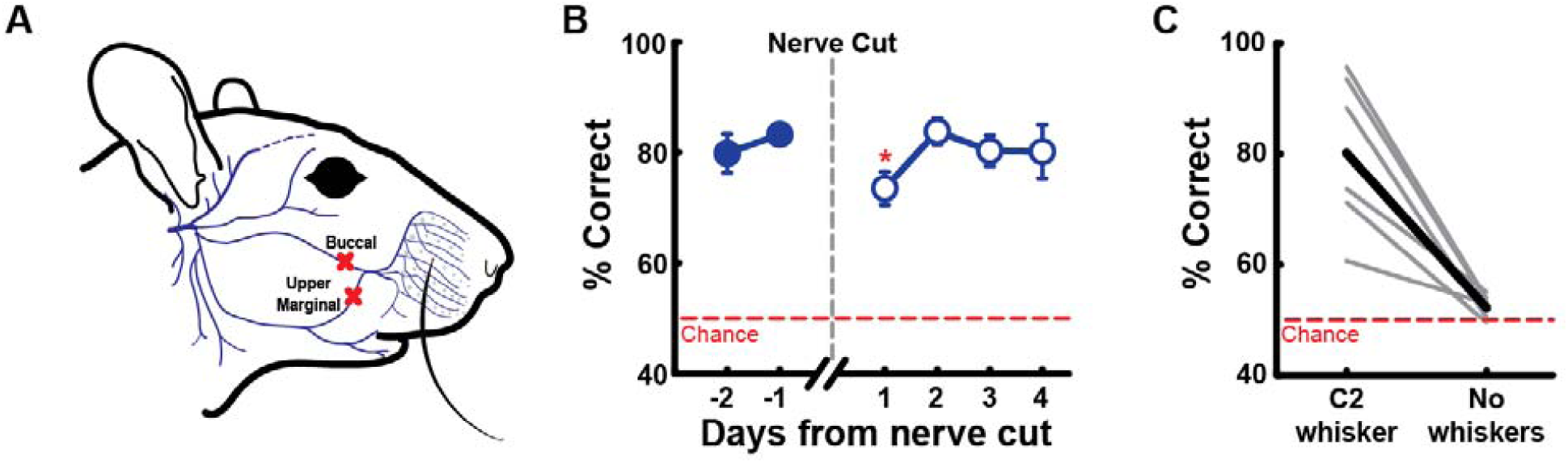
Mice do not require whisker motor control to discriminate passively presented textures. **(A)** Facial motor nerve lesions targeting buccal and upper marginal to immobilize whisker pad. Adapted from (Heaton et al 2014). **(B)** Behavioral performance before and after facial motor nerve lesion. Mice can discriminate between passively presented smooth-rough textures in the absence of whisker motor control (n = 7 mice). Data shown as mean±SEM. \**P*<0.05 compared to pre-nerve cut day -1. **(C)** Post-facial nerve lesion whisker trimming confirms that task remained whisker-dependent (average in black, individual mice in gray, n = 7 mice).

In summary, these results demonstrate that active movement is not necessary for the discrimination of textures, at least in the range of texture differences in our design. This also demonstrates that the effect of barrel cortex lesion on this behavior involves sensory deficits and cannot reflect the lesion disrupting only movement.

### Barrel cortex is dispensable for 2AFC object detection

Prior work from our lab showed S1 to be dispensable for an active, whisker-dependent go/no-go pole detection task (Hong et al 2018). Are these divergent outcomes due to essential computations mandated by the task (texture discrimination versus object detection) or to the different task designs (2AFC versus go/no-go)? We considered the possibility that the 2AFC task design is itself cortex dependent. We therefore developed a 2AFC object detection task identical to that used for discrimination except animals were presented with either a surface (textures from Table 1) or no surface (Fig 4A). Mice quickly learned a passive version of the task, with 100% reaching learning criterion within nine days in the assessment phase (4.0 ± 0.5 d, range 2-9 d; Fig 4B,C). They also readily detected objects with a single whisker (Fig 4D, Video 6), with performance recovering to full-whisker levels (see pre-lesion Fig 4E) typically within 3-4 days. For a separate active cohort of mice, the object was moved further away so that the animals had to actively whisk forward to detect the objects (Video 7).

**Figure 4.**
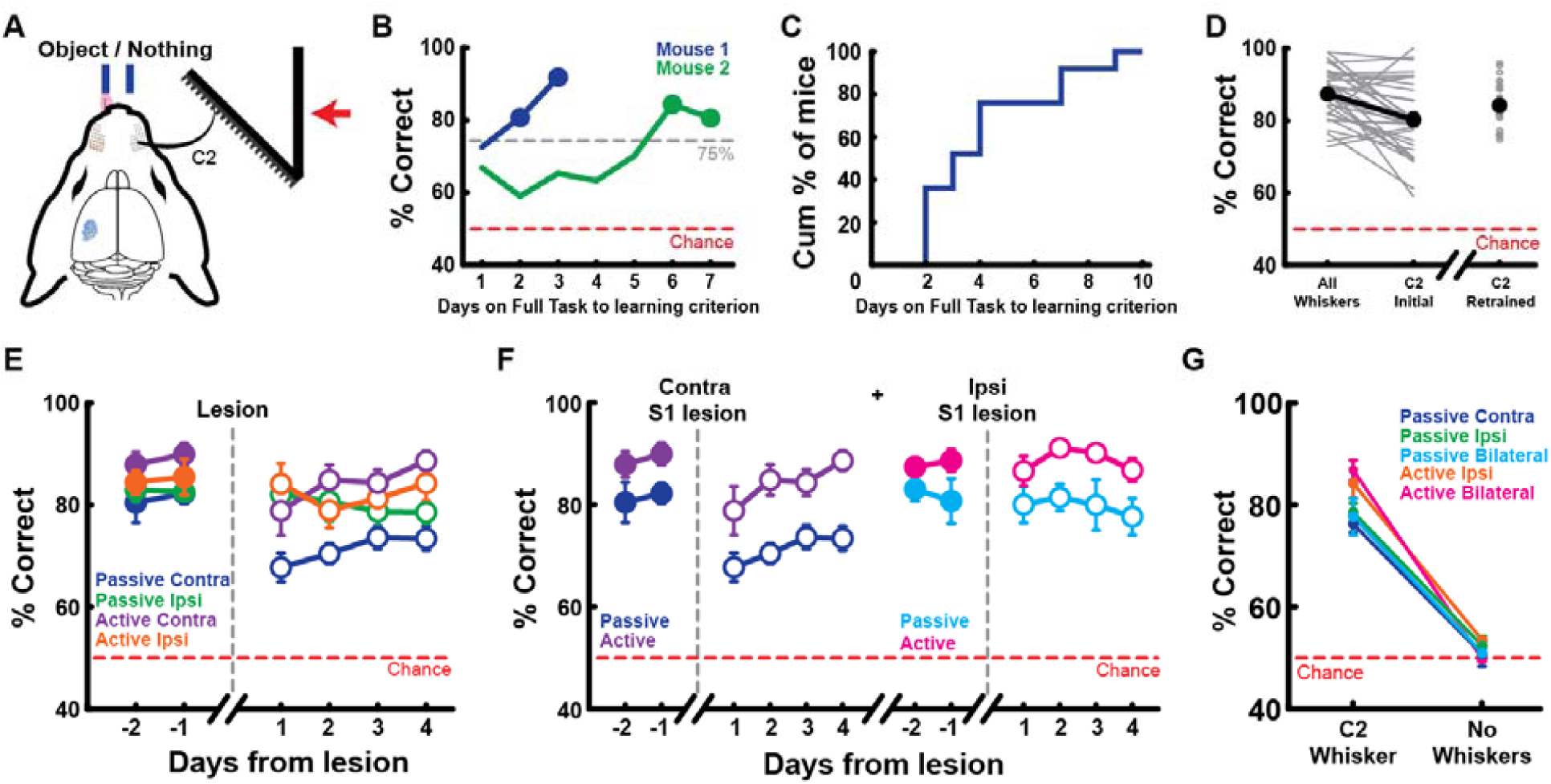
Barrel cortex is dispensable for a 2AFC object detection task. **(A)** Head-fixed two-alternative forced choice object detection paradigm. Mice learned to lick left for an object (textures from Table 1), and lick right for nothing. **(B)** Example learning curves of mice reaching learning criterion (≥75% correct on two consecutive learning assessment days). **(C)** Cumulative histogram of number of days on Full Task to reach learning criterion (n = 25/25 mice). **(D)** Mice readily detect objects using a single whisker (average in red, individual mice in gray, n = 32 mice). Behavioral performance typically recovers to all-whisker levels and are ready for manipulations within 3-4 days. **(E)** Lesioning contralateral barrel cortex minimally impairs performances on both passive and active 2AFC detection (passive contralateral n = 8 mice, passive ipsilateral n = 6 mice, active contralateral n = 6 mice, active ipsilateral n = 6 mice). Task performance eventually recovers to pre-lesion levels by day 7 (extended recovery not shown). **(F)** Ipsilateral barrel cortex does not compensate for loss of contralateral barrel cortex on either the passive or active 2AFC detection tasks (passive contralateral n = 8 mice, active contralateral n = 6 mice, passive ipsilateral n = 4 mice, active ipsilateral n = 6 mice). **(G)** Whisker trim control confirms all detection behaviors are whisker-dependent (passive contralateral n = 4 mice, passive ipsilateral n = 6 mice, passive bilateral n = 4 mice, active ipsilateral n = 6 mice, active bilateral n = 6 mice). Data shown as mean±SEM.

For both passive and active cohorts, once mouse performance stabilized, we aspirated the barrel cortex as described above to test its involvement in 2AFC object detection. Behavior was partially impaired one day after lesioning the contralateral barrel cortex (passive *P*<0.001; active *P*=0.015; paired *t*-test) but still remained significantly above chance (50%; *P*<0.001, one-sample *t*-test; Fig 4E, purple and blue). Performance gradually recovered to pre-lesion levels by Day 4 and 7 for active and passive detection, respectively (extended recovery not shown, see pre-Ipsi lesion in Fig 4F). While the post-lesion performance of the passive group (67.7%) was lower overall than the active group (78.8%), these groups also had different baseline pre-lesion performance (passive 82.2%, active 90.0%) and the same relative effect of the lesion (compare blue and purple): we observed no differences when comparing passive and active performance deficits as a proportion to pre-lesion baseline (*P*=0.29, two-sample *t*-test). Control ipsilateral barrel cortex lesions did not impair performance for either passive or active cohorts (passive *P*=0.85; active *P*=0.68; paired *t*-test; Fig 4E, green and orange).

In the whisker system, there are no connections across the midline linking left and right trigeminal brainstem complexes or thalamic nuclei (Frangeul et al 2014). Nevertheless, to be absolutely certain that ipsilateral barrel cortex did not compensate for loss of the contralateral, we ablated the ipsilateral barrel cortex in a subset of mice with contralateral lesions. We found no performance impairments (Fig 4F), demonstrating that an intact ipsilateral cortex does not compensate for contralateral lesions on this task. Together, these results demonstrate that barrel cortex plays a key role during our texture discrimination task, not because of its 2AFC design but rather because barrel cortex is essential to texture-related computations.

Given prior disparate outcomes of transient S1 inactivation, we also examined the effects of optogenetic manipulations on the 2AFC object detection task. The Emx1-Halo mouse line (Supplementary Fig 3A) permits direct, selective, and robust silencing of excitatory cortical neurons across all layers (Hong et al 2018). After training Emx1-Halo mice to perform passive 2AFC object detection with a C2 whisker, we located the corresponding C2 barrel via intrinsic signal optical imaging. Optogenetically silencing barrel cortex with 40 mW of laser power impaired overall behavior (reduced from 82.3% to 60.0% averaged across a minimum of four days; *P*<0.001, paired *t*-test). Although the optogenetically induced impairment was more substantial than when we lesioned barrel cortex, behavior again remained significantly above chance (50%; *P*<0.001, one-sample *t*-test; Supplementary Fig 2B, green). Control Flex-Halo mice lacking Cre were behaviorally unaffected by the laser (from 89.1% to 88.1%; *P*=0.57, paired *t*-test; Supplementary Fig 2B, black).

In a go/no-go object detection paradigm, optogenetic S1-silencing reduces overall responses to both go and no-go trials (Hong 2018), leaving the possibility that behavioral deficits could be due to more general impairments in the ability to respond. In our 2AFC detection paradigm, where animals are required to respond to both conditions, animals had similar response times during laser ON or OFF trials (ON/OFF response comparison between Emx1-Halo and Flex-Halo, *P*=0.55; two-sample *t*-test; Supplementary Fig 3C), suggesting that optogenetic inactivation did not impair the response behavior, per se. Indeed, Emx1-Halo mice tended to report that nothing was present during optogenetic silencing (Emx1-Halo *P*=0.022; Flex-Halo *P*=0.47; one-sample *t*-test; Supplementary Fig 3F,G). This “no object” response bias reflected a causal effect of barrel cortex inactivation rather than a byproduct of task uncertainty; animals showed no such bias when their performance was impaired from trimming down to a single whisker (Supplementary Fig 3H, see Fig 4D for whisker trim impairment). Animals showed increased response for “no object” when all their whiskers were trimmed (Supplementary Fig 3H). The whisker trim controls also reduced detection task performance to chance, confirming that mice did not overcome lesions or optogenetic silencing by exploiting other sensory modalities (Fig 4G, Supplementary Fig 3E).

These results demonstrate that detection can occur in the absence of barrel cortex, even when mice are required to perform in more sophisticated 2AFC design. In summary, barrel cortex is essential for texture discrimination, and task design (2AFC versus go/no-go) is not a critical determinant of cortical dependence.

### Transient cortical inactivation impairs texture discrimination

While dispensable for object detection, cortex is clearly necessary for texture discrimination as demonstrated by our chronic manipulations. Transient optogenetic manipulations therefore provide a distinct opportunity to test the functional contribution of specific cortical layers. We first tested Emx1-Halo animals (Fig 5A) to characterize the impact of inactivating all cortical layers. Mice were trained on the passive smooth-rough task. Transient inactivation of all cortical layers reduced performance fully to chance (reduced from 75.7% to 49.1%; *P*<0.001, paired *t*-test; Fig 5B, green), consistent with our lesion manipulation. Control Flex-Halo animals were unaffected by the laser (83.2% to 83.4%; *P*=0.57, paired *t*-test; Fig 5B, black). Response time did not differ between genotypes (ON/OFF response comparison, *P*=0.40; two-sample *t*-test; Supplementary Fig 5A), consistent with cortical inactivation not affecting licking motor response. Whisker trimming confirmed that mice did not use other sensory modalities (Fig 5C).

**Figure 5.**
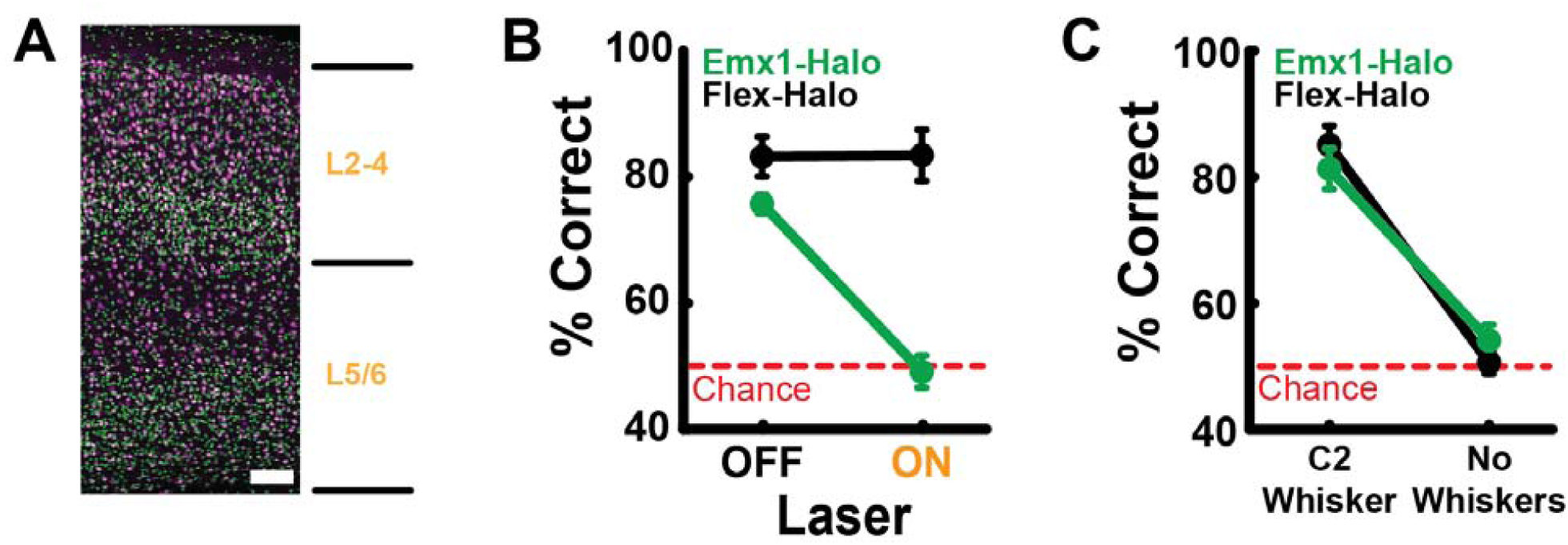
Transient barrel cortex inactivation impairs texture discrimination to chance. **(A)** Coronal section of Emx1-GFP mouse cortex. Green, GFP; magenta, NeuN; scale bar, 100 μm. **(B)** Behavioral performance for laser OFF versus ON trials (Emx1-Halo n = 4 mice, Flex-halo n = 4 mice). Optogenetic inactivation of barrel cortex impairs smooth-rough texture discrimination performance to chance. **(C)** Whisker trim control confirms that task remained whisker-dependent. Data shown as mean±SEM.

Interestingly, we observed learning deficits in Emx1-Halo animals. Most non-learners were Emx1-Halo mice (of 76 mice that underwent smooth-rough task training, 14/17 non-learners were Emx1-Halo). Only 8/22 Emx1-Halo mice (36%; Supplementary Fig 4A, green) learned the passive smooth-rough discrimination task compared to 7/7 wildtype (100%; *P*=0.0033, Chi-Square; black) and 44/47 of all other transgenic animals on the wildtype background (93%; *P*<0.001, Chi-Square; orange). Of the learners, Emx1-Halo mice were also slower to reach criteria (15.5 ± 2.5 d, mean±SEM) compared to wildtype (8.0 ± 1.6 d, mean±SEM; *P*=0.028, two-sample *t*-test) and other transgenic animals (8.5 ± 0.9 d, mean±SEM; *P*=0.0031, two-sample *t*-test). However, despite these learning deficits, Emx1-Halo animals that did learn the task performed similarly to other genotypes.

We noticed that for several Emx1-Halo mice (excluded from Fig 5), optogenetic silencing with 40 mW laser power resulted in lower performance even during laser OFF trials: six Emx1-Halo mice exhibited inconsistent and low performance on laser OFF trials (see Methods, Supplementary Fig 4C,D, green). This effect was not observed during our 2AFC detection (Supplementary Fig 3) or go/no-go task (Hong et al 2018) likely because detection was cortex independent whereas texture discrimination was cortex dependent. The laser OFF deficit did not occur when light-proof black tape was placed between optic fiber and cortex (*P*=0.63, two-sample *t*-test; Supplementary Fig 4B), consistent with cortical inactivation having a lingering effect rather than laser luminance startling the animals. Reduced power (20 mW) did not disrupt laser OFF performance while still reducing laser ON performance to chance (Supplementary Fig 4D, cyan).

In summary, transient cortical inactivation impaired texture discrimination performance to chance. Combined with the lesion results, the two manipulations demonstrate that sensory discrimination of tactile textures requires the barrel cortex.

### Deep and superficial layers are critical for texture discrimination

Having firmly established through both chronic and transient manipulations that texture discrimination requires barrel cortex, we next examined the roles of the deep and superficial cortical layers. Since thalamocortical inputs are sufficient to drive activity in both sets of layers separately, it is possible that upper and deep layers could function independently of one another depending on the task. To test whether either the superficial or deep layers suffice to perform the behavior, we trained transgenic lines that target Cre and therefore halorhodopsin to various combinations of cortical layers. We again used the single-whisker passive smooth-rough texture discrimination task to remove any requirement for whisking. None of the layer-specific transgenic lines displayed learning deficits (Supplementary Fig 4A, orange) or sensitivity to high laser power. Hence, for consistency, 40 mW laser power was used for the remaining experiments.

The deep cortical layers receive direct thalamic input and contain major cortical outputs to subcortical structures relevant for behavior. We targeted excitatory pyramidal neurons in the deep layers by crossing a transgenic mouse line expressing Cre recombinase mainly in L5 but also to some extent in L6 (Rbp4-Cre) (Gerfen et al 2013) with the stop-Halo reporter (Fig 6Ai). Texture discrimination of trained Rbp4-Halo animals was severely impaired by photoinhibition (reduced from 84.8% to 58.7%; *P*<0.001, paired *t*-test; Fig 6Aii).

**Figure 6.**
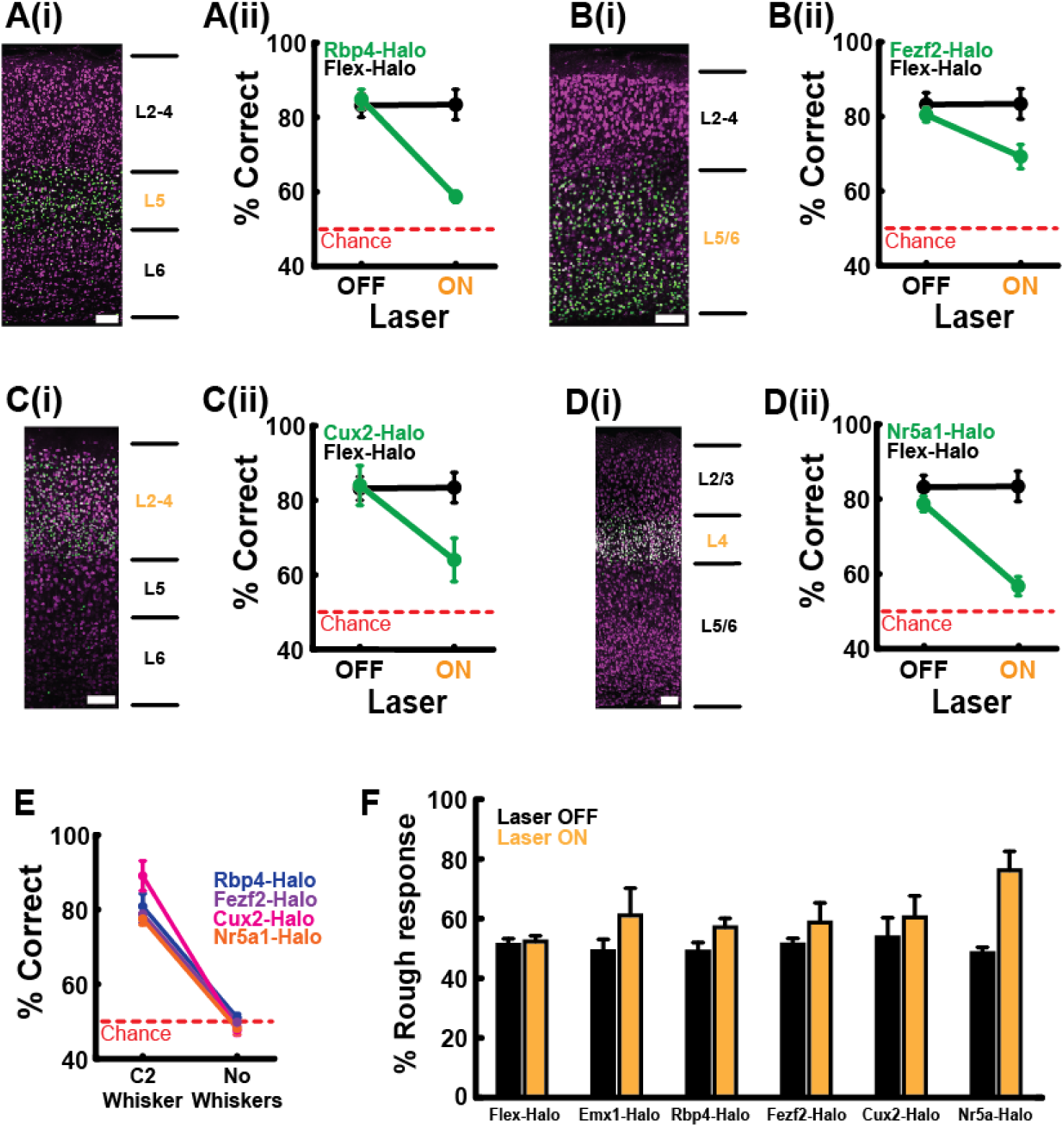
Deep and superficial layers are critical for texture discrimination. **(A-D)** Effects of layer-specific inactivation on smooth-rough texture discrimination performance. **(i)** Green, GFP; magenta, NeuN; scale bar, 100 μm. **(ii)** Behavioral performance for laser OFF versus ON trials. Flex-Halo (n = 4 mice). **(A)** Rbp4-Halo (n = 5 mice). **(B)** Fezf2-Halo (n = 7 mice). **(C)** Cux2-Halo (n = 3 mice). **(D)** Nr5a1-Halo (n = 5 mice). **(E)** Whisker trim control confirms all smooth-rough discrimination behaviors are whisker-dependent. **(F)** Silencing any cortical layers increased response selectivity for rough textures. Flex-Halo *P*=0.17; Emx1-Halo *P*=0.16; Rbp4-Halo *P*=0.039; Fezf2-Halo *P*=0.12; Cux2-Halo *P*=0.14; Nr5a-Halo P=0.0072 compared to 50%. Data shown as mean±SEM.

Because the deep layers contain distinct cell types that project to different downstream targets, we tested an additional deep layer transgenic mouse line. We crossed the stop-Halo reporter line with Fezf2-CreERT2 animals expressing inducible recombinase. After tamoxifen administration to adult animals (P>120) to activate halorhodopsin expression, we observed inhibitory opsin in L5 pyramidal neurons and sparsely in L6 (Fig 6Bi). Optogenetic manipulations significantly impaired behavioral performance (reduced from 80.4% to 69.2%; *P*=0.0011, paired *t*-test), but not to the same extent as in Rbp4-Halo mice (ON/OFF, *P*=0.0011, two-sample *t*-test; Fig 6Bii).

Rbp4-Halo and Fezf2-Halo experiments demonstrated that deep cortical layers are needed for texture discrimination. However, the main anatomical target of thalamocortical axons is L4, which in turn innervates L2/3 (Petersen 2019). Additionally, others have previously shown that a subset of long-range cortico-cortical projecting S1 neurons in L2/3 are recruited during texture discrimination but not object detection (Chen et al 2013). Therefore, to test the role of the excitatory L2-4 neurons in this superficial pathway, we crossed Cux2-CreERT2 mice (Franco et al 2011) with stop-Halo animals (Fig 6Ci). As with the deep layers, photoinhibition of the superficial layers impaired performance of Cux2-Halo mice but only at a trend level (reduced from 83.7% to 63.1%; *P*=0.11, paired *t*-test; Fig 6Cii).

L4 receives an order of magnitude more thalamocortical synapses than L2/3 does (Oberlaender et al 2012). Within L4, the barrels are innervated by the ventral posterior medial nucleus (VPM) of the thalamus while the septal neurons between barrels are modulated by higher-order posterior medial nucleus (POm) (Bruno & Sakmann 2006, Bruno & Simons 2002, Wimmer et al 2010). Because the POm-septal pathway is far less responsive to sensory stimuli than the VPM-barrel pathway (Diamond et al 1992), we used transgenic mice expressing Cre recombinase in excitatory L4 barrel neurons (Nr5a1-Cre) but not the septal neurons that lie between the barrels (Fig 6Di). Nr5a1-Halo animals were severely impaired by photoinhibition (reduced from 78.7% to 56.7%; *P*<0.001, paired *t*-test; Fig 6Dii), demonstrating that the pathway through the barrels is critical for texture discrimination. Together, the Cux2ERT2 and Nr5a1 data show that the superficial layers via VPM→L4→L2/3 are as important to texture discrimination as the deep layer VPM→L5/6 pathway.

The response times for all tested transgenic lines were comparable to Flex-Halo (ON/OFF response comparisons for Rbp4-Halo *P*=0.89, Fezf2-Halo *P*=0.31, Cux2-Halo *P*=0.84, Nr5a-Halo *P*=0.89; two-sample *t*-tests; Supplementary Fig 5B-E). Thus, behavioral impairments upon inactivating subsets of layers are due to sensory rather than global motor deficits. At the end of the experiments, whisker trimming of all tested lines confirmed that mice did not switch to using other sensory modalities (Fig 6E).

L2/3 has a strongly suppressive effect on deep layer activity (Adesnik & Scanziani 2010, Pluta et al 2019). Conceivably, strong connections like these could produce local forms of diaschisis, whereby silencing one set of layers does not reveal its task involvement but rather that it disrupts another key set of layers. We therefore investigated whether layer-specific manipulations produced different response biases. If our results were due merely to local diaschisis, one might expect opposing effects of deep and superficial layer silencing. On the contrary, silencing any layer increased the likelihood of animals responding as if texture was rough (Fig 6F, Supplementary Fig 6A-E). This rough response bias is observed in each and every Cre line. The response bias cannot be due to simple motor bias (lick direction) because the same manipulation caused animals performing our detection task to lick the opposite direction (Supplementary Fig 3G). Additionally, the increased likelihood of animals responding to textures as rough reflect a causal effect of barrel cortex inactivation rather than a confound of animal uncertainty since trimming down to a single whisker – which impaired task performance (Fig 2B, blue) – did not lead to a response bias (Supplementary Fig 6F). Similarly, response biases were not observed when all whiskers were cut off (Supplementary Figure 6F). These analyses indicate that unintended local circuit disruption effects do not explain the important role of deep and superficial layers in our task.

Taken together, our results show that, despite independently receiving direct sensory input from the thalamus, both the deep and superficial layers are critical for texture perception.

## Discussion

Despite extensive, detailed studies of cortical microcircuitry, the computations performed by cortical layers during sensory processing and their functional contributions to behavior remain unclear. Investigating these issues requires a cortically dependent behavior. We developed a novel 2AFC whisker-based texture discrimination task for head-fixed mice. After barrel cortex lesion, trained mice could not perform this task again, demonstrating that texture discrimination is cortex dependent. This stands in stark contrast to our past and present results regarding object detection. Interestingly, whisker-paralyzed animals could discriminate textures without active whisking. Using layer-specific optogenetic manipulations, we discovered that both the deep and superficial layers of cortex are similarly involved in this behavior. Thus, our work establishes an experimentally tractable, sensory-guided behavior for which both the deep and superficial layers are essential.

### Parametric 2AFC texture discrimination task to study cortical function

Understanding how cortical layers contribute to sensory-guided behaviors requires a thorough characterization of a cortex-dependent task. Hence, we developed a texture discrimination behavior that (1) uses parametrically controlled surfaces, (2) utilizes a sophisticated 2AFC design, (3) is amenable to high-throughput training, (4) facilitates completeness of neural manipulations, and (5) requires the cortex.

We controlled the roughness of our textures by changing the width, depth, and spacing between the grooves; this allows quantification of psychometric curves for whisker-mediated texture discrimination. Indeed, we showed that animals immediately generalize to novel textures using this approach (Fig 1). This differs from many prior studies using sandpaper stimuli that vary greatly between manufacturers and cannot be easily parameterized (Oladazimi et al 2018).

Additionally, our 2AFC task design provides more rigorous analysis of behavior compared to the go/no-go task design that is a staple of mouse sensory behavioral studies. Go/no-go designs are susceptible to fluctuations in animal motivation (Shenoy & Angela 2012), whereby subjects show increased bias towards go responses, leading to higher hits and false alarm rates. For this reason, nonhuman primate studies have used 2AFC task design for decades. Only recently have human experimentalists identified choice reporting methods and shaping procedures that allow training of rodent subjects on head-fixed 2AFC tasks (Guo et al 2014a). We find mice rapidly — sometimes in as little as a few days — learn our 2AFC task design in both texture discrimination and object detection behaviors.

Our single-whisker paradigm is ideal for future physiology studies, allowing focusing of neural recordings on a single cortical column. We can also ensure the completeness of our neural manipulations by targeting the corresponding column and calibrating our effect to include nearby columns. Ideally manipulations must include these neighboring columns because the thalamic neurons often have surround receptive fields that include adjacent whiskers (e.g., a C2 barreloid neuron may respond to C1, C2, and C3) (Brumberg et al 1996, Bruno & Simons 2002).

Additionally, texture discrimination involves numerous other structures beyond primary somatosensory cortex, such as secondary somatosensory cortex, motor cortex, association cortex, prefrontal cortex, striatum, and cerebellum (Chen et al 2013, Gilad et al 2018, Manita et al 2015, Proville et al 2014, Simões-Franklin et al 2011). For example, the magnitude of texture roughness is first thought to be processed in S1 subsequently followed by feature extraction to detect roughness changes in the higher-order secondary somatosensory cortex (Jiang et al 1997). Therefore, our 2AFC texture discrimination task is an attractive paradigm for future studies investigating the role of specific genes, cell types, or nervous system structures in normal behavior and disease.

### What does sensory cortex do?

Sensory cortex may have evolved to achieve one or more functions. The longest standing idea is that sensory cortex exists to construct progressively more complicated feature detectors (Gilbert & Wiesel 1979, Marr & Thach 1991). In this scenario, cortical circuits establish selectivity for features that do not exist at the level of individual primary afferents. Sensory cortex has also been proposed to relate sensory inputs to models of the body and its environmental context (Brecht et al 1997), functioning more like a traditionally conceived association cortex. For instance, classic lesion studies found that barrel cortex is unnecessary for detection, but is required in the context of a freely-moving gap cross behavior in which the animal must detect a location of a platform relative to its head and body (Hutson & Masterton 1986). Because our textures were presented at a consistent spatial location relative to the animals, our results demonstrate that both feature detection and egocentric spatial mapping functions may coexist in barrel cortex.

In contrast to sensory features, an object’s presence or absence is readily decoded from even a single primary afferent during detection behaviors. The functional role of primary sensory cortex in detection has, however, been controversial across visual, auditory, gustatory, and somatosensory behaviors (Feldmeyer et al 2013, Stüttgen & Schwarz 2018). Different paradigms, task difficulty, and inactivation methods have yielded conflicting results. Transient pharmacological and optogenetic S1 inhibition impaired object tactile detection behaviors (Guo et al 2014b, Hong et al 2018, Miyashita & Feldman 2013, O’Connor et al 2010, Sachidhanandam et al 2013). However, the sudden loss of S1 may have disrupted downstream areas that are vital for detection behavior, such as the striatum (Otchy et al 2015, Wolff & Ölveczky 2018). Indeed, when S1 but not striatum is lesioned, mice could still learn and execute a whisker-dependent go/no-go object detection task (Hong et al 2018). Our own 2AFC detection results demonstrate that, even in a different paradigm, the cortex is not required for detection behaviors in which neural responses to the two sensory conditions (object/no object) are trivial to classify by any downstream structure. This indicates that cortex is dispensable for head-fixed sensory detection, irrespective of task design.

Interestingly, the recovery rates of our active 2AFC objection detection animals were similar to our previous active go/no-go detection animals (Hong et al 2018) while passive 2AFC mice recovered more slowly. A likely explanation for this difference is that passive 2AFC mice are receiving stimuli closer to sensory threshold, which likely places greater demands on neural circuits. Whiskers are moving at nearly 2,000°/sec during active objective detection (Hong et al 2018) whereas passively presented objects are advanced into the whiskers at a much lower velocity. The subcortical circuitry that could mediate detection after lesion may be less sensitive to some feature of these passive stimuli, such as their low velocity. Cortex is indeed known to be tuned for high velocity events (Khatri et al 2004) whereas subcortical structures such as brainstem and thalamus may be less so or tuned to entirely different features (Minnery & Simons 2003, Pinto et al 2000).

Here, we clearly demonstrate that texture discrimination requires barrel cortex. Only one previous study has ever causally tested the role of barrel cortex in texture discrimination (Guic-Robles et al 1992). In that study, freely moving rats were trained to distinguish between two textured platforms in order to jump across a gap to the reward-associated platform. Because that design utilized a gap cross paradigm similar to the freely moving detection tasks discussed above (Hutson & Masterton 1986), texture may not have been the key feature rendering the task cortex dependent, but rather the need to integrate within the body and environment maps. Our head-fixed design eliminated such spatial components and directly tested representation of texture in sensory cortex. Thus, barrel cortex is essential to representing this tactile quality.

Multiple features of a surface are thought to contribute to the perception of texture (Lederman 1974, Lederman 1983). The exact nature of the sensory cues that differentiate textures, however, remains an open question. Various hypotheses have been proposed for the whisker system, analogous to the skin of the fingertip interacting with surfaces. First, sequences of friction that trap the whisker followed by high velocity “slips” (“stick-slip” events) have been shown to vary in rate with sandpaper and gratings of various roughness (Isett et al 2018, Jadhav et al 2009). In this scenario, velocity tuning might then explain the cortex dependency of texture discrimination. Second, tuning of whiskers to different frequencies as whiskers resonate following slips has been hypothesized to play a role in texture encoding (Neimark et al 2003). Resonance seems an unlikely mechanism given that our mice were minimally impaired when their whisker inputs were reduced to a single whisker (i.e., C1-C2-C3 → C2). Furthermore, it is unclear why cortex would be required in this scenario as resonance is a property of the whiskers and could already be decoded at the level of the primary afferents. Future comparisons of cortical and subcortical encoding of features that covary with surface roughness may reveal why cortex is so essential in texture discrimination.

Stick-slip events have previously been shown to increase with roughness and drive cortical activity (Jadhav et al 2009). Optogenetic inhibition should decrease cortical activity that would normally reflect such stick-slip events, rendering the animals’ perception of surfaces as smooth. On the contrary, our mice tended to report all textures as rough when we inactivated all or any set of cortical layers. These results seem to run counter to the notion of stick-slip encoding, at least in our particular paradigm. An intriguing possibility is that, instead of stick-slip or resonance encoding, spectral features of whisker microvibrations may carry substantial amounts of texture information, a possibility under investigation in the non-human primate somatosensory system (Lieber & Bensmaia 2019, Weber et al 2013).

### Does texture discrimination benefit from active movement?

Active movement is suspected to enhance sensory discrimination (Gamzu & Ahissar 2001, Gibson 1962, Kleinfeld et al 2006). Passive and active sensory discrimination fundamentally differ not only at the behavioral level but also at the neural level. For example, Krupa et al (2004) demonstrated increases in both the duration and magnitude of S1 activity evoked by active discrimination compared to relatively brief and weaker transient activity during passive discrimination. Moreover, they found increased duration and degree of response suppression during passive compared to active discrimination. Increased post-central gyrus, prefrontal, and cerebellum activity for active discrimination has also been reported in humans (Simões-Franklin et al 2011). Hypothetically, active exploration might therefore engage higher order processes at both the behavioral and neural levels to enhance discrimination.

Surprisingly, our mice discriminated passively and actively presented textures equally well. By making active movements during passive presentations, mice could have confounded our design. However, paralyzing the whisker pad, which guaranteed that the task was executed in a truly passive form, did not prevent mice from discriminating the tested texture pairs. Performance on passive and active versions of the same texture discrimination task has rarely been directly compared previously. A few psychophysical studies that matched passive presentation speeds to active movements found human performance on passive discrimination rivaled active discrimination (Lederman 1981, Lederman 1983), consistent with our results using a moderately difficult task. A similar comparison in an fMRI study found increased S1 activity during active compared to passive discrimination of simple textures despite no differences in behavioral performance (Simões-Franklin et al 2011). Further studies are needed to test whether active sensation only becomes beneficial for the finer, more challenging discriminations.

### Emx1-Halo animals have abnormal learning on a cortically dependent task

We found that Emx1-Halo transgenic mice struggle to learn the cortex-dependent 2AFC texture discrimination task. Layer-specific lines did not exhibit learning deficits. Once fully trained, Emx1-Halo mice performed comparably to other genotypes. Interestingly, we did not observe Emx1-Halo learning deficits on the detection task, likely because cortex is dispensable for learning object detection (Hong et al 2018). Hence, the Emx1-Halo learning abnormality is likely unique to cortex-dependent tasks.

Induced opsin expression can adversely affect cell health through toxic or leaky expression and cause neural and behavioral abnormalities (Allen et al 2015). Driving calcium indicators with an Emx1-Cre line can induce epileptiform events, which may result from the combination of calcium indicator overexpression and Cre toxicity (Steinmetz et al 2017). Analogously, opsin overexpression and Cre-expression induced abnormalities (Kim et al 2013) may jointly contribute to the learning deficits we observed in Emx1-Halo mice. Emx1-Cre mice without responder transgenes have never before been tested on a task equivalent to ours. Future studies are needed to determine if Cre toxicity alone in such an abundant cortical population is sufficient to impair learning.

### Both deep and superficial layers are essential for texture discrimination

Both the deep and superficial layers of cortex receive direct sensory input from the thalamus (Constantinople & Bruno 2013). This raises the intriguing possibility that the two sets of cortical layers constitute different processing systems. What are possible purposes of separate networks? One possibility is that the deep and superficial layers are redundant networks that support cortex-dependent behaviors. If so, either set of layers should be sufficient for the behaviors. Our results demonstrate that, despite receiving the same sensory input, all layers are critical for our texture discrimination task, ruling out redundancy of cortical layer function.

A second scenario is that deep and superficial layers could have unique, independent functions, compatible with their known physiology and anatomy. Their constituent cell types differ in intrinsic membrane, synaptic integration, and spike generation properties (Palmer et al 2014, Zhao et al 2016). Moreover, both sets of layers in barrel cortex receive inputs from diverse sources (primary thalamus, secondary thalamic nuclei, motor cortex, and secondary somatosensory cortex), but only deep layer neurons have substantial innervation by secondary thalamus and cortex in the vicinity of their basal dendrites (Wimmer et al 2010, Zhang & Bruno 2019). Additionally, superficial layers almost exclusively target other cortical areas with the exception of a small population of corticostriatal neurons. The deep layers, however, target both cortical and subcortical structures including behaviorally relevant areas (striatum, colliculus, brainstem). The results of our layer-specific manipulations are compatible with unique, independent functions (e.g., both superficial and deep layer projections to other systems are necessary).

A third scenario, however, is that the tightly interconnected layers are synergistic. For instance, simple sensory information could be processed directly via thalamocortical projections to cortical output neurons in L5/L6, while more complex top-down sensory computations require feedback from the superficial layers (Adesnik & Naka 2018, Constantinople & Bruno 2013, Petersen 2019). At a general level, our data are also compatible with this hypothesis. However, the most prominent synergy observed to date showed L2/3 suppresses deep layer activity (Adesnik & Scanziani 2010, Pluta et al 2019), predicting opposite results of deep versus superficial manipulations. We find on the contrary that mice tend to classify textures as rough when either deep or superficial layers are inhibited. This lends support to the notion of unique, independent functions, but further experiments – perhaps manipulating connections between the layers or between a layer and its long-range target – are needed to definitively distinguish these two hypotheses. It also conceivable that whether the layers have unique, independent functions versus synergistic interaction depends heavily on the specific task the animal engages in at that moment.

---

**Supplementary Figure 1.**
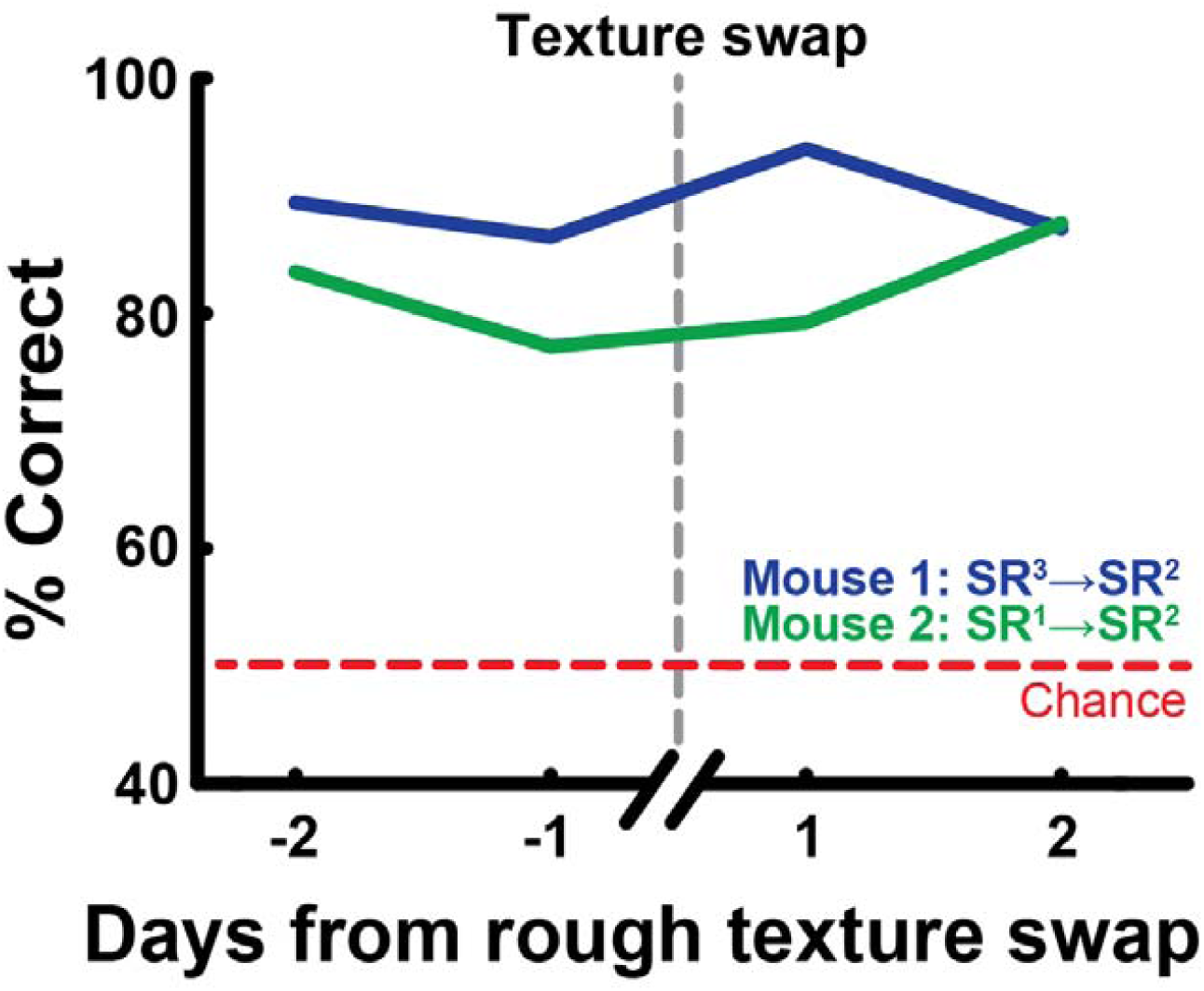
Mice readily generalize to discriminate rough textures. Example performance of two mice generalizing to discriminate smooth from a novel rough texture. See Table 1 for quantification of R^1^, R^2^, and R^3^ rough textures used in the smooth-rough (SR) discrimination paradigm.

**Supplementary Figure 2.**
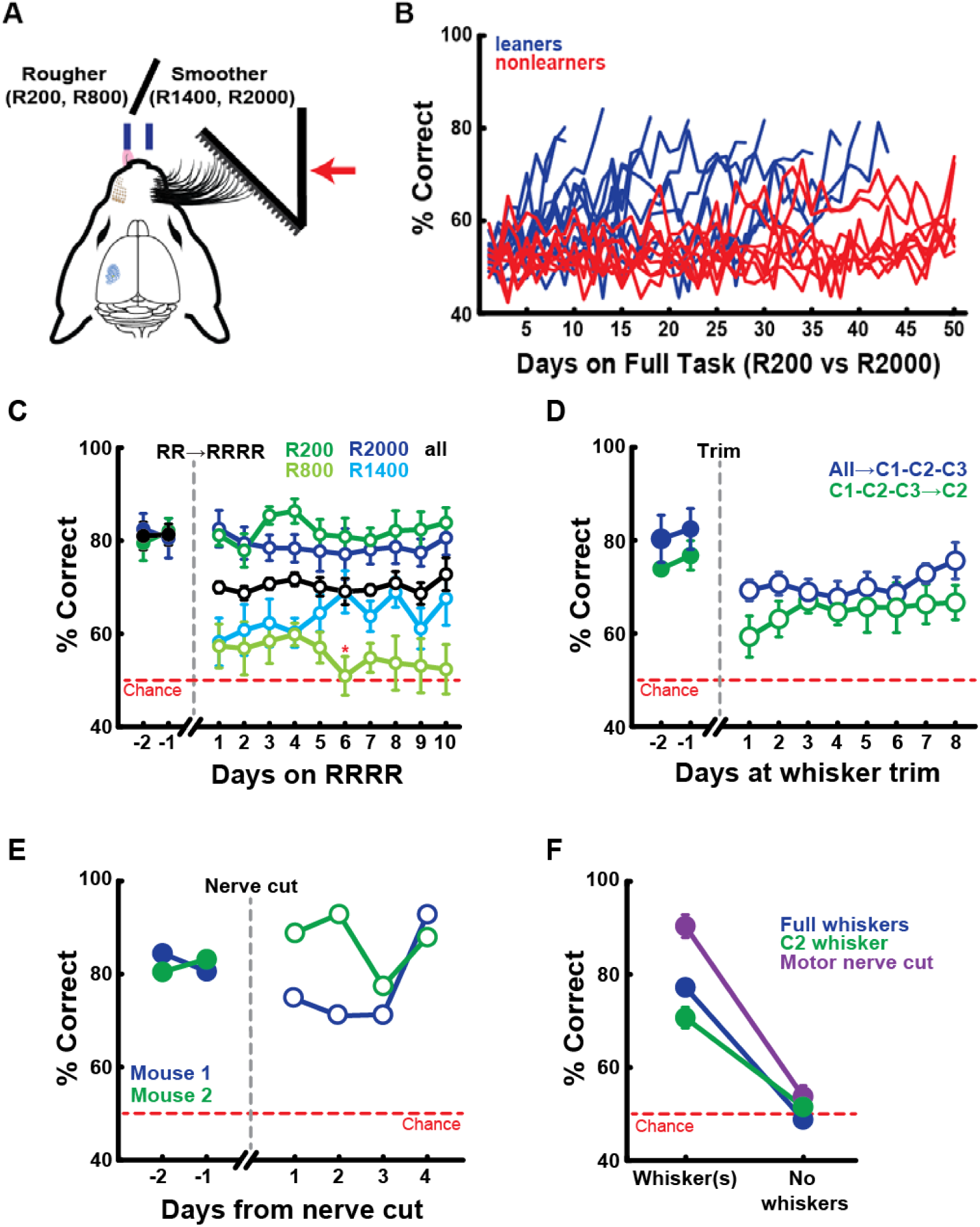
Mice can discriminate rough textures. **(A)** 2AFC paradigm for rough-rough (RR) and four texture discrimination (RRRR). See Table 2 for texture quantifications. **(B)** Learning curves of mice reaching learning criterion on rough-rough paradigm (≥75% correct on two consecutive learning assessment sessions within 50 Smart Random days, learners n = 11, nonlearners n = 9). **(C)** Behavioral performance when exposed to all four rough textures (days -2 to 7 n = 9 mice, days 8 to 10 n = 8 mice). \**P*<0.05 compared to R1400 performance. **(D)** Mice can discriminate rough-rough textures using three (n = 4 mice) or even a single whisker (n=4 mice, 3/4 mice reached consistent +70% performance at single whisker by day 8, individual mouse data not shown). **(E)** Fully-whiskered mice can discriminate rough-rough textures without whisker motor movement (n = 2 mice). **(F)** Whisker trim controls confirm that rough-rough discrimination with full whiskers (n = 6 mice), C2 whisker (n = 3 mice), and motor nerve cut (n = 2 mice) are whisker-dependent. Data shown as mean±SEM.

**Supplementary Figure 3.**
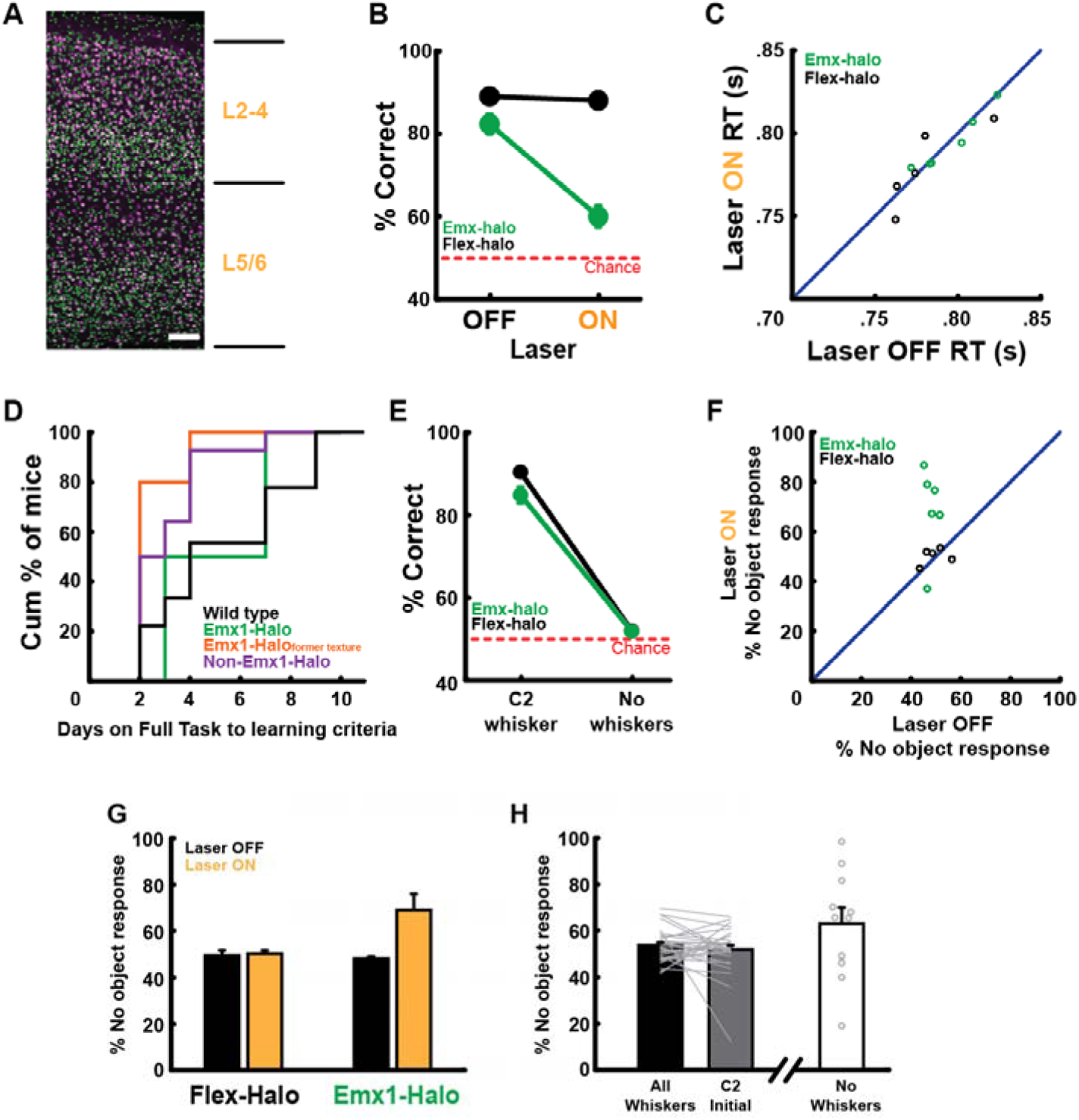
Transient photoinhibition of cortex impairs object detection and increases likelihood of animals reporting “no object”. **(A)** Coronal section of Emx1-GFP mouse cortex. Green, GFP; magenta, NeuN; scale bar, 100 μm. **(B)** Behavioral performance for laser OFF versus ON trials (Emx1-Halo n = 6 mice, Flex-Halo n = 5 mice, each mouse performance averaged across 4+ sessions). Optogenetic inactivation of barrel cortex impairs detection behavior. **(C)** No differences in response latency between genotypes and laser ON/OFF trials (Emx1-halo n = 6 mice, flex-halo n = 5 mice, each mouse reaction time averaged across 4+ sessions). Response time calculated as duration of time between the approximate first whisker contact and the first committed lick that corresponds to the choice lick during the response window. **(D)** Cumulative histogram showing number of days on Full Task to reach learning criterion. Wildtype n = 9 mice, Emx1-Halo mice n = 2 mice, Emx1-Halo mice who were non-learners on the texture discrimination task n = 5 mice, Non-Emx1-Halo transgenic mice n = 14 mice. Emx1-Halo animals formerly on texture discrimination were not included in the detection learning curve analysis. **(E)** Whisker trim control confirms that the detection behaviors are whisker-dependent. **(F**,**G)** Emx1-Halo mice show increased response selectivity for “no object” during laser ON trials. **(F)** Silencing barrel cortex in Emx1-Halo mice biased 5/6 animals to respond more as if no object is present. **(G)** Genotype averages for response selectivity. Laser ON: Flex-Halo *P*=0.47, Emx1-Halo *P*=0.022 compared to 50%. **(H)** Increasing uncertainty and impairing task performance by trimming down to a single whisker did not alter likelihood of mice reporting “no object”. *P*=0.33 compared to 50% (n = 30 mice). Trimming the last whisker marginally increased preference for reporting “no object”. *P*=0.092 compared to 50% (n = 11 mice). Data shown as mean±SEM.

**Supplementary Figure 4.**
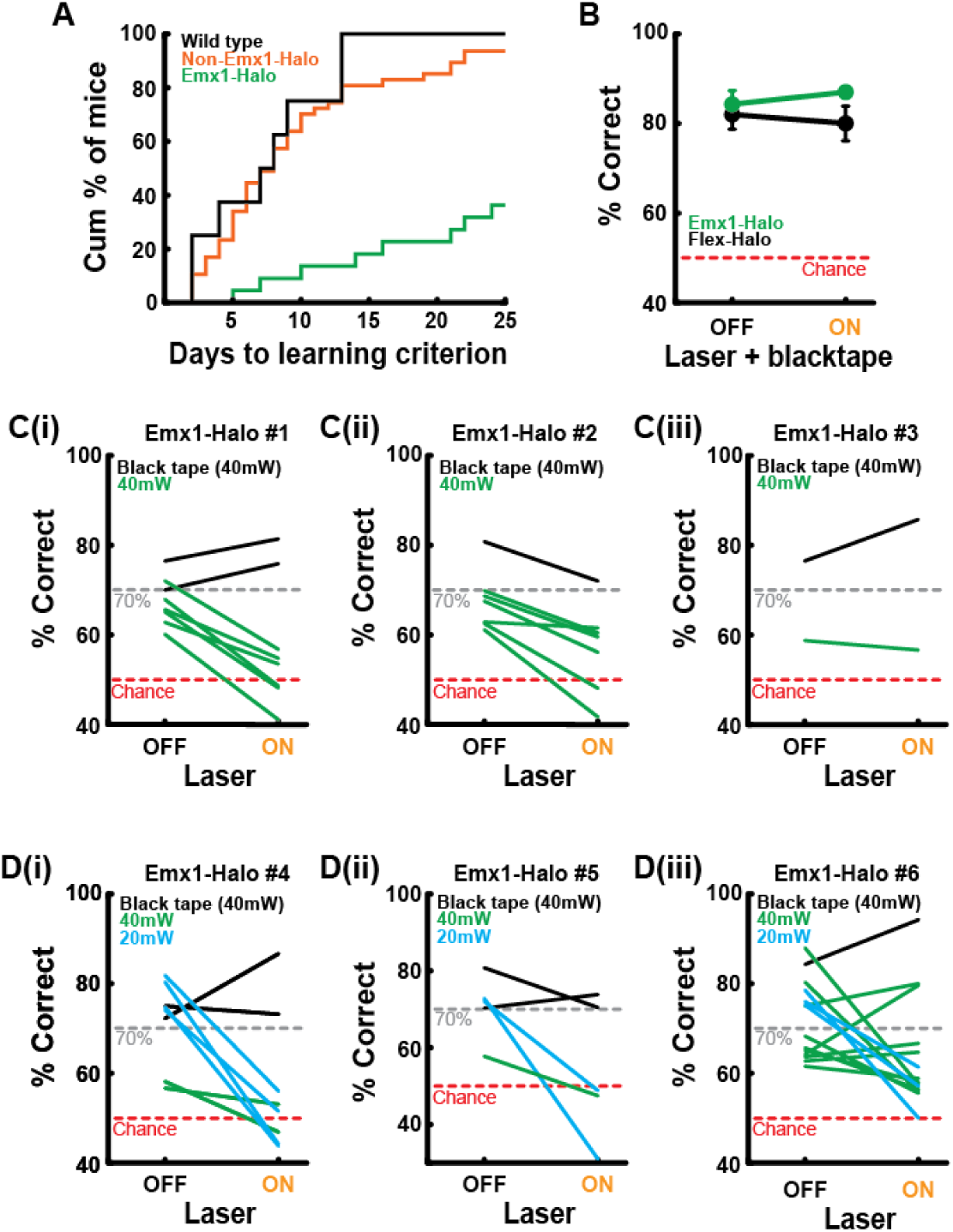
Emx1-Halo mice exhibit learning impairments and are sensitive to transient cortical inactivation. **(A)** Cumulative histogram showing number of days on Smart Random to reach learning criterion. Wildtype n = 7 mice, Non-Emx1-Halo transgenic mice n = 47 mice, Emx1-Halo mice n = 22 mice. **(B)** Control black tape optogenetic sessions show no behavioral impairment on neither laser OFF or ON trials for both Emx1-Halo (n = 4) and Flex-Halo (n = 4) mice. **(C-D)** Emx1-Halo animals are sensitive to transient cortical inactivation. **(C)** Control black tape sessions (black) confirm that physically blocking the laser does not affect behavioral performance. Emx1-Halo mice are sensitive to the 40 mW laser (green), with their performance on laser OFF trials often falling below the 70% cut-off criteria. **(D)** When the laser power is decreased to 20 mW (cyan), Emx1-Halo animals are able to perform above the 70% criteria on OFF trials. We still observe at-chance performance during laser ON trials. Data shown as mean±SEM.

**Supplementary Figure 5.**
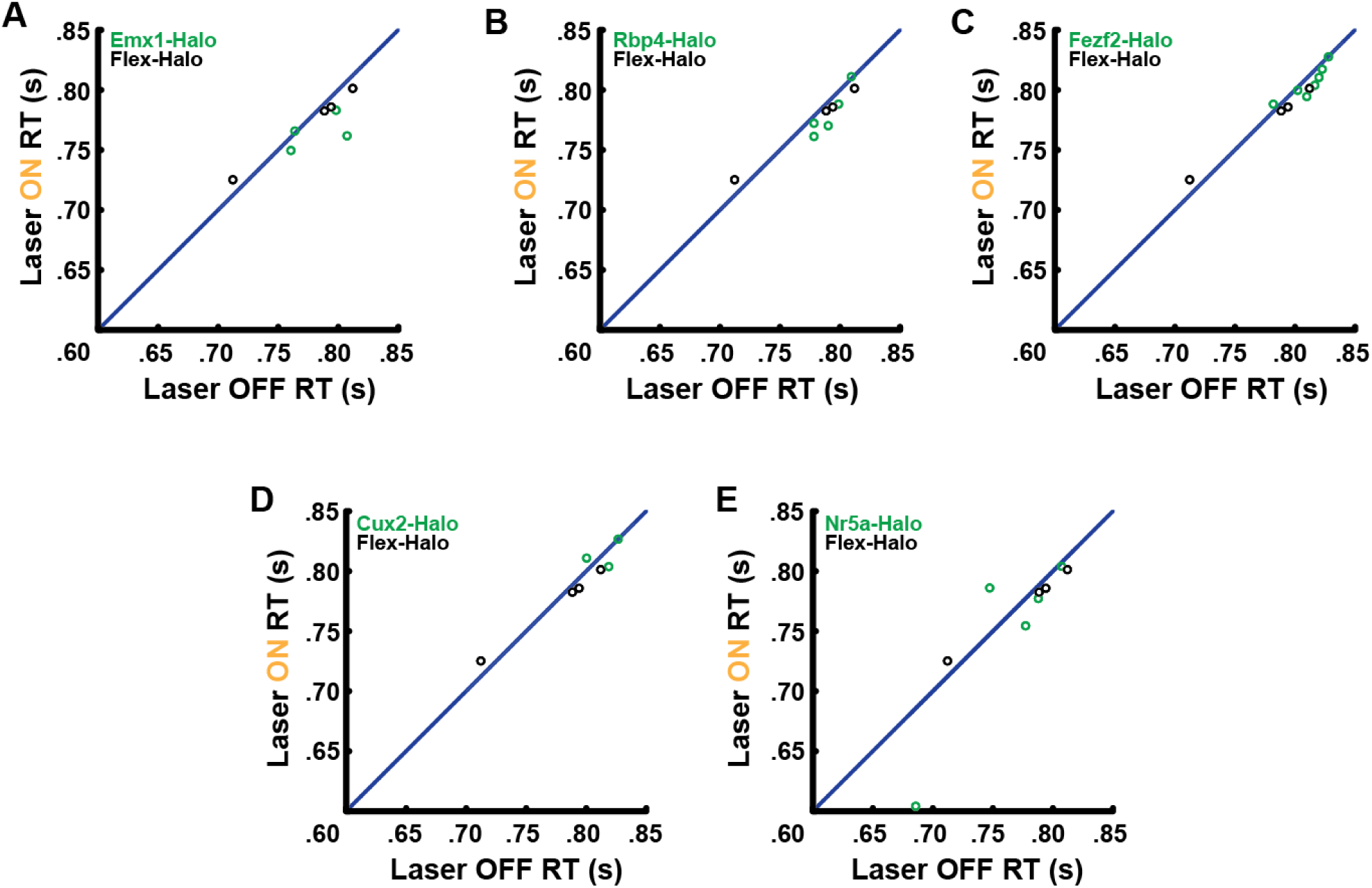
Cortical photoinhibition does not affect response latencies. **(A-E)** Response time for laser OFF versus ON trials, calculated as duration of time between the approximate first whisker contact and the first committed lick that corresponds to the choice lick during the response window. Flex-Halo (n=4 mice), black. **(A)** Emx1-Halo (n=4 mice). **(B)** Rbp4-Halo (n=5 mice). **(C)** Fezf2-Halo (n=7 mice). **(D)** Cux2-Halo (n=3 mice). **(E)** Nr5a-Halo (n=5 mice).

**Supplementary Figure 6.**
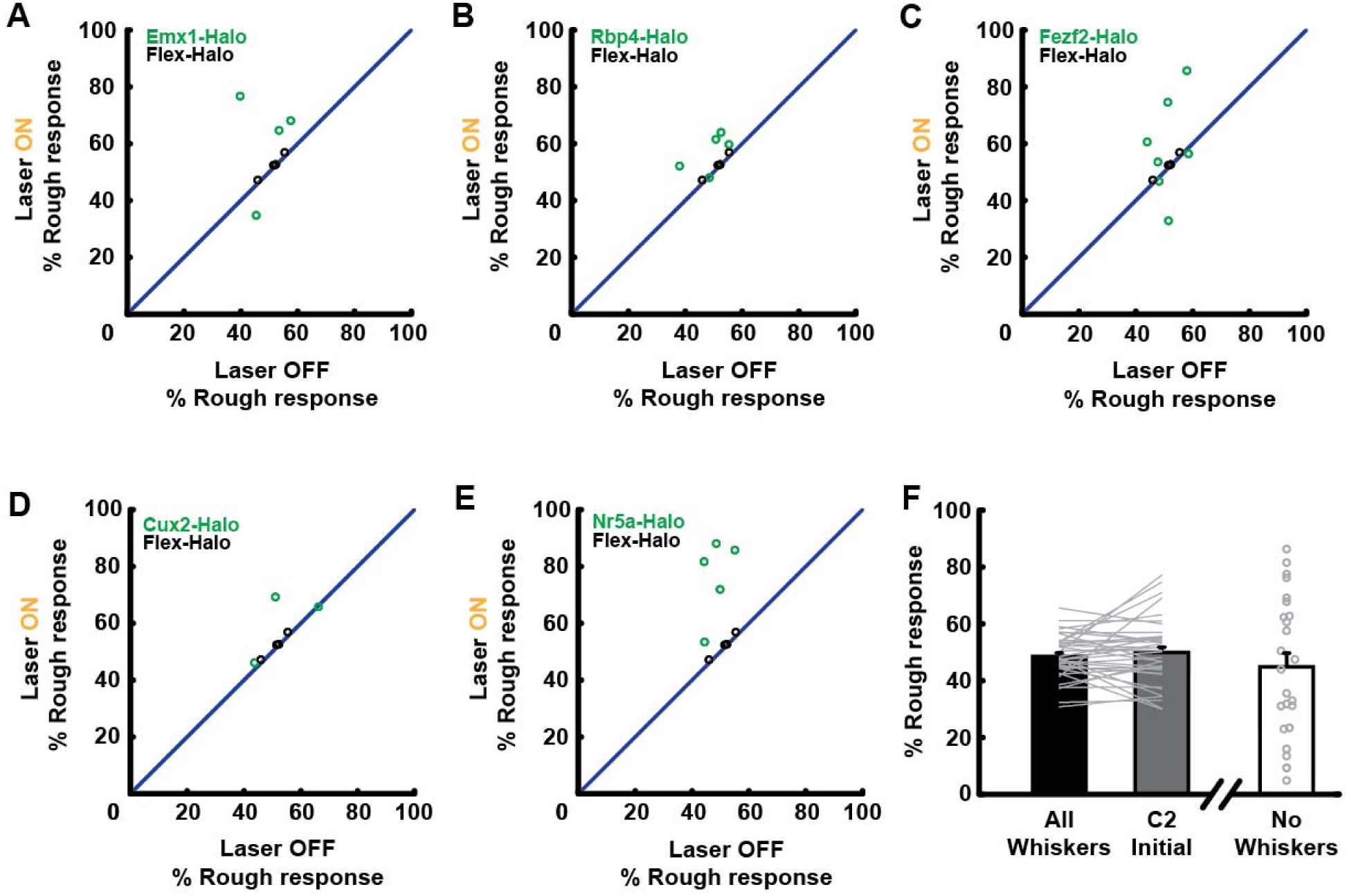
Cortical photoinhibition increases response bias for rough textures. **(A-E)** Silencing any cortical layers biased mice to respond more as if the rough texture is presented. Flex-Halo (n = 4 mice), black. **(A)** Emx1-Halo (n = 4 mice). **(B)** Rbp4-Halo (n = 5 mice). **(C)** Fezf2-Halo (n = 7 mice). **(D)** Cux2-Halo (n = 3 mice). **(E)** Nr5a-Halo (n = 5 mice). **(F)** Animals did not exhibit preference for reporting textures as rough when trimmed to a single whisker (*P*=0.98 compared to 50%, n = 39 mice) or no whiskers (*P*=0.30 compared to 50%, n = 26). Data shown as mean±SEM.

## STAR Methods

### Resource Availability

#### Materials Availability

This study did not generate any unique reagents.

#### Data and Code Availability

The datasets and code generated during this study will become publicly available upon publication of this manuscript.

### Mice

Experiments involving live animals complied with the NIH Guide for the Care and Use of Laboratory Animals and were approved by the Institutional Animal Care and Use Committee at Columbia University. All mice used in experiments were adults (>P120) and were maintained on a 12:12 light/dark cycle with *ad libitum* access to food and water. *Ad libitum* access to water was removed for two days before behavioral training and testing. Access to food was never restricted. Mice were maintained at ∼80% of their free-drinking weight. All data were collected from mice of either sex.

We report data from 35 object detection (13 male, 22 female), 91 smooth-rough texture discrimination (47 male, 44 female), and 20 rough-rough texture discrimination (11 male, 9 female) animals. 10/35 detection animals and 15/91 smooth-rough mice were excluded from learning analysis due to changes in behavioral setup, lab relocation, or extended break in training due to trainers’ schedules. Except for the rough-rough discrimination mice with C57BL/6J; 129S2/SvPasCrl background, all mouse lines were maintained on a C57BL/6 background from Jackson Laboratories.

Emx1-Halo mice were generated by crossing Emx1-IRES-Cre knock-in mice (Jackson Laboratories, stock #005628) to Rosa-lox-stop-lox (RSL)-eNpHR3.0/eYFP mice (stop-Halo Ai39, JAX, stock# 006364), as previously described (Hong et al 2018). Rbp4-Halo, Fezf2-Halo, Cux2-Halo, and Nr5a1-Halo mice were generated by crossing Rbp4-Cre-KL100, Fezf2-CreER, Cux2-CreERT2, and Nr5a1-Cre with the stop-Halo animals and were heterozygous for indicated transgenes. Cre-negative Flex-Halo control mice did not express the halorhodopsin transgene. The experimenters were blind to the genotype of the C57BL/6J; 129S2/SvPasCrl rough-rough discrimination animals for an ongoing experiment. However, the experimenters were not blind to the genotype of all other animals during testing and analysis.

### Method Details

#### Tamoxifen Injections

Adult Cux2-Halo and Fezf2-Halo mice (>P120) carried inducible recombinase Cre-ERT and were intraperitoneally injected with tamoxifen (20 μL of a 20 μM/μL solution in corn oil) to induce Cre expression. These animals were injected five times every other day for optimal Cre expression.

#### Headplate surgery

Adult mice (>P120) were anesthetized using isoflurane and administered buprenorphine and carprofen. Under a stereotaxic setup, we removed the scalp and implanted a custom-designed stainless steel headplate (Wilke Enginuity) over the skull with Metabond dental cement (Parkell).

#### Behavioral setup

We adapted a 2AFC shape discrimination task (Rodgers et al 2020) to implement the 2AFC texture discrimination and object detection tasks. The behavioral apparatus was controlled by a microcontroller (Arduino Uno) and housed in a box (Foremost) custom-designed with a lightproof door and covers. The apparatus was built on an aluminum bread board (Thorlabs or Newport), posts and bases (Thorlabs), and custom-designed laser-cut acrylic pieces. In the discrimination task, a linear actuator (Actuonix L12-30-50-6-R) advanced laser-cut acrylic textures (see Tables 1 and 2) towards the right whiskers of the animal. The textures were passively presented at 45° relative to the animal’s midline, with the surfaces gently pushing into the majority of whiskers (∼1 cm from the right whisker pad, see Video 1). The response window opened soon after the textures reached their final position.

The mouse indicated its choices by licking two stainless steel pipes (McMaster) that were positioned slightly anterior and to either side of its mouth. While the animals pre-emptively licked before the response window opened, only the first lick within the window determined whether the trial was correct or incorrect. Response latency was calculated as duration of time between the approximate first whisker contact and the first committed lick that corresponds to the choice lick during the response window. The animals were trained to lick right for a smoother surface and to lick left for a rougher surface (Fig. 1A, Supplementary Fig 2A) for a water reward (∼5 μL). The water was delivered by breaking an infrared beam (Sparkfun, QRD1114) or triggering capacitive touch sensors (Sparkfun MPR121) that opened solenoid valves (The Lee Co. LFAA1209512H). The linear actuator then moved the object sufficiently far from the mouse to prohibit any whisker contact, and a stepper motor (Pololu 1204) rotated the object wheel to a neutral position before rotating to the next trial’s object. Rotation to a neutral position prior to object presentation ensured that the sound and vibration generated by the motor was consistent between trials and no distinct auditory or vibrational cues were given. Additionally, a white light (LE LED; Amazon B00YMNS4YA) was flashed in-between trials to prevent mice from fully dark-adapting and using visual cues to perform the task. A computer fan (Cooler Master; Amazon B005C31GIA) also blew air over the texture and away from the mouse to prevent the animals from picking up olfactory cues. The textures were also regularly cleaned with 70% ethanol.

The detection task setup was identical to that used for discrimination except that the linear actuator either presented a surface (textures from Table 1) or no surface (Fig 4A).

#### Animal training

After 1 week recovery, headplated and water-restricted mice (>P120) were put through the following training protocol (Fig 1C).

1. Handling and freely moving lick training. At this stage, animals were handled by trainers for ∼4 days and placed in the behavioral apparatus where they could roam and habituate to their environment. The mice also had access to the water lick pipes from which they could obtain water.
2. Head-fixed lick training. Mice were head-fixed and initially taught to obtain water from a single lick pipe. They were then required to lick left and right lick pipes in an alternating pattern, rotating between the two after a set number of licks which progressively decreased to 1, in which case the reward-side would switch between every lick. This stage required ∼5 days of training.
3. Forced Alternation. Here, texture stimuli were presented and associated with a reward lick pipe, and a punishment time-out was introduced. Presentation of the same texture was repeated until mice responded correctly, after which the other texture was presented. The timeout was initially 2 s and gradually increased to 9 s as performance increased. Trials in which no responses were recorded within the response window (45 s) were excluded from analysis. Detection, smooth-rough discrimination, and rough-rough discrimination animals required 6.8 ± 0.4, 12.0 ± 0.6 and 14.9 ± 5.0 (mean±SEM) days respectively.
4. Full task. Each day started with 45 forced alternation trials to ease the animals into the full task. After these initial trials, randomized left/right trials were presented, and licks continuously monitored for biases. When a bias was detected, the program halted the randomized trials and adjusted to counteract the bias. For example, if the animal responded ≥20% more on one side than the other, the program delivered continuously delivered trials corresponding to the other side. If the mouse developed a perseverative stay bias (*P*<0.05, ANOVA), the program delivered forced alternation trials. Only the complete randomized trials were used to assess performance. Learning criterion was defined as ≥75% correct performance for 2 consecutive days. Detection, smooth-rough discrimination, and rough-rough discrimination mice reached criterion in 9.2 ± 0.8, 25.0 ± 3.5, and 4.0 ± 0.5 (mean±SEM) days respectively.
5. Whisker trimming. Once mice reached learning criterion, all whiskers except C2 whisker were trimmed off under anesthesia. Detection animals transitioned from all to one whisker (Fig 4D). Texture discriminating mice either experienced the same all→C2 or more gradual all→C1-C2-C3→C2 protocol (Fig 2B, Supplementary Fig 2D). Trimming was maintained regularly for the remainder of the experiments. However, a subset of mice substantially struggled after whisker trimming and required their trimmed whiskers to grow back in order to regain mastery of the behavior. At this point, animals were advanced to completely random sessions unless the experimenter manually corrected for an unusual side bias. At the end of the experiments, animals that did not fall to chance (50%; cutoff ≤60%) after full whisker trimming (all including C2) were excluded from the study.
6. Active task. For a subset of both the detection and the smooth-rough discriminating animals, the final stimulus position was gradually positioned further away until the mice had to actively whisk forward with their remaining whisker to contact the textures. These animals often required additional training to master these new circumstances.

#### Intrinsic signal optical imaging

Animals were anesthetized with isoflurane, and the skull over the left barrel cortex (centered ∼1.5 mm posterior of bregma and 3.5 mm lateral of the midline) was drilled thin over a 4–5 mm area. A thin layer of superglue was applied over the thinned skull. Images were acquired with a CCD camera (Q-Imaging, Rolera-MGi Plus) mounted on a stereomicroscope and software custom-written in LabVIEW. The vasculature on the brain surface was imaged with 510/40 band-pass filtered illumination (Chroma, D510/40), and intrinsic signal imaging done with illumination with a 590-nm long-pass filter (Thorlabs, OG590). The C2 whisker was stimulated with 10 pulses at 5 Hz with a multi-directional piezo stimulator in the dorsal/ventral directions. The location of the maximum reflectance change was mapped relative to the surface vasculature. Surrounding whiskers were also imaged to confirm the identification of the C2 barrel.

#### Cortical lesions

Mice were anesthetized under isoflurane. A 3-4 mm craniotomy was made over the target C2 barrel. The underlying cortical tissue was aspirated with a sterile blunt-tipped syringe connected to a vacuum. Lesions encompassed the C2 column and its immediately adjacent columns as well as all cortical layers. The majority of lesions extended beyond the barrel cortex boundaries into the body subdivision of S1 (Fig 2C). Lesioned animals were allowed to recover for 1 day after surgery prior to testing. After completing the task, mice were perfused, and the brains extracted for histological analysis. Brains were coronally sliced (100 µm thick) with a vibratome. All lesions were objectively scored by experienced anatomists who were blind to the behavioral data. Lesions that were scored to have extended beyond S1 and into subcortical areas like the striatum were excluded from analysis.

#### Videography

During training in the behavioral apparatus, animals were monitored using a Sony PlayStation 3 Eye Camera under infrared illumination (Video 1). For high-speed videography, we moved the behavioral setup into a customized lightproof box mounted on a vibration-isolating air table (TMC). Fully trained single-whisker mice were recorded using a high-speed camera (Photonfocus DR1-D1312IE-100-G2-8 and Basler ace acA720-520um) under infrared illumination (Videos 2-7).

#### Optogenetic inactivation

Optogenetic experiments were performed in the lightproof box mounted on the vibration-isolating air table (TMC). All optogenetic animals performed passive versions of their respective tasks (detection or smooth-rough discrimination) with a single whisker. A 593-nm laser (OEM) was turned on for 25% of the trials, which were randomly interleaved with laser-off trials. The laser was coupled to a 200-µm diameter, 0.39 NA optic fiber (Thorlabs) via a fiberport. The fiber tip was positioned over C2 cortical column using the vasculature-referenced intrinsic signal map. For laser-on trials, an optical shutter opened at the onset of the trial as the linear actuator advanced the textures towards the mouse. The texture moved within reach of the whisker field ∼1s after the onset of the laser, ensuring photoinactivation before contacts were possible. The laser remained on until the mouse responded, or, if no response was made, for a maximum of 3 s. Trials in which the response occurred after the 3 s were excluded from analysis. No tissue damage was detectable with the described protocol.

#### Confocal imaging

Mice were perfused, and the brains coronally sectioned (100 µm thick) on a freezing microtome. Slices were stained with a rabbit α-NeuN (Millipore MAB5406) and mouse α-GAD67 (Millipore ABN78) primary antibodies (1:100, incubated for 3-4 days) followed by anti-rabbit conjugated to Alexa Fluor 555 (Life Technologies) and anti-mouse conjugated to DyLight 633 (Invitrogen) secondary antibodies (1:500, overnight). Slices were imaged on a Nikon A1R confocal microscope using a 20x/0.75 NA air objective. Imaging depth ranged 20-30 µm at 5 µm intervals.

### Quantification and statistical analysis

#### Statistics

Analyses were done with custom-written scripts in MATLAB and Python. For all figures, statistical significance is denoted as \**P*<0.05, \*\**P*< 0.01, \*\*\**P*< 0.001. Differences were not significant (*P*>0.05) unless otherwise indicated.

## Acknowledgements

We thank Drew Baughman for assistance with histology and genotyping; Lindsay Jordan for assistance with histology; Esther Greeman and Nina Harano for assistance with mouse training; Caroline Bollen for assistance with microscopy, and Ramin Khajeh for advice on data analysis.

Funding was provided by NIH R01 NS094659 (RMB); NSF GRFP and T32NS064928 (JMP); NIH F32 NS084768 and T32 MH015174 (YKH); and NIH F32 NS096819, Kavli Institute Postdoctoral Fellowship, and Brain & Behavior Research Foundation Young Investigator Award (CCR).

